# Resolution doubling in light-sheet microscopy via oblique plane structured illumination

**DOI:** 10.1101/2022.05.19.492671

**Authors:** Bingying Chen, Bo-Jui Chang, Philippe Roudot, Felix Zhou, Etai Sapoznik, Madeleine Marlar-Pavey, James B. Hayes, Peter T. Brown, Chih-Wei Zeng, Talley Lambert, Jonathan R. Friedman, Chun-Li Zhang, Dylan T. Burnette, Douglas P. Shepherd, Kevin M. Dean, Reto P. Fiolka

**Affiliations:** Lyda Hill Department of Bioinformatics, University of Texas Southwestern Medical Center, Dallas, TX, USA; Cecil H. and Ida Green Center for Systems Biology, University of Texas Southwestern Medical Center, Dallas, TX, USA; Department of Cell Biology, University of Texas Southwestern Medical Center, Dallas, TX, USA; Aix Marseille University, CNRS, Centrale Marseille, I2M, Turing Centre for Living Systems, Marseille, France; Department of Cell and Developmental Biology, Vanderbilt Medical Center, University of Vanderbilt, Nashville, TN, USA; Center for Biological Physics and Department of Physics, Arizona State University, Tempe, AZ, USA; Department of Molecular Biology, University of Texas Southwestern Medical Center, Dallas, TX, USA; Department of Cell Biology, Harvard Medical School, Boston, MA, USA; Department of Systems Biology, Harvard Medical School, Boston, MA, USA

## Abstract

Structured illumination microscopy (SIM) doubles the spatial resolution of a fluorescence microscope without requiring high laser powers or specialized fluorophores. However, the excitation of out-of-focus fluorescence can accelerate photobleaching and phototoxicity. In contrast, light-sheet fluorescence microscopy (LSFM) largely avoids exciting out-of-focus fluorescence, thereby enabling volumetric imaging with low photo-bleaching and intrinsic optical sectioning. Combining SIM with LSFM would enable gentle 3D imaging at doubled resolution. However, multiple orientations of the illumination pattern, which are needed for isotropic resolution doubling in SIM, are challenging to implement in a light-sheet format. Here we show that multidirectional structured illumination can be implemented in oblique plane microscopy, a LSFM technique that uses a single objective for excitation and detection, in a straightforward manner. We demonstrate isotropic lateral resolution below 150nm, combined with lower photo-toxicity compared to traditional SIM systems and volumetric acquisition speed exceeding 1Hz.

## Main Text

In fluorescence microscopy, one must carefully balance spatiotemporal resolution, imaging depth, and the duration of time one can image prior to the onset of photobleaching or phototoxicity^1^. For example, super-resolution microscopy improves the spatial resolution of an imaging system well-beyond the diffraction limit, but in most cases requires thin specimens, high laser powers, and the acquisition of many imaging frames (or independent measurements) and is therefore often slow and phototoxic. In contrast, by adopting a unique imaging geometry that reduces the illumination burden placed on a specimen, Light-Sheet Fluorescence Microscopy (LSFM) has evolved in an orthogonal direction and improved imaging speed while reducing phototoxicity, albeit at diffraction-limited resolutions^2-5^.

For years, researchers have sought to advantageously combine the strengths of LSFM with super-resolution microscopy and thereby enable gentle, longitudinal, and volumetric imaging of molecular processes at sub-diffraction spatial scales^6^. Using localization-based super-resolution methods that rely on stochastic transitions between non-radiative and radiative states (e.g., Photoactivation Localization Microscopy^7^ and Stochastic Optical Reconstruction Microscopy^8^), several LSFM implementations have been reported^9-11^ Nonetheless, owing to the limited photon collection efficiency of the used objectives, or lossy polarization-based detection paths, deleteriously high laser powers were necessary to achieve single-molecule detection^12, 13^. In contrast, by launching the illumination beam with a reflective cantilever or microfluidic device, high numerical aperture objectives can be leveraged^14-17^. Nonetheless, live cell imaging remains challenging, as single molecule localization approaches require long acquisition times (depending on the labeling density, reaching in the most extreme case up to days^18^).

Alternatively, by leveraging methods such as Stimulated Emission Depletion^19^ which selectively suppress fluorescence from molecules residing outside of a region of interest, axial resolution well below the diffraction limit has been realized in LSFM^20, 21^. While stimulated emission is a general mechanism compatible with many fluorophores, the involved high laser power densities are opposite to what LSFM tries to achieve, e.g., lowering the irradiation burden placed upon the sample. This has in part been addressed by using photo-switching of fluorophores instead of stimulated emission to improve the axial resolution^22^, at the cost of restricting the choice of usable fluorophores. Nonetheless, neither method has so far been leveraged to improve the lateral resolution in LSFM.

Structured Illumination Microscopy (SIM)^23^, while not achieving the same spatial resolution as localization based approaches or stimulated emission depletion, features high temporal resolution and works with any fluorophore in low to medium laser power regimes (on the order of 1-10W/cm^2^). While a combination of SIM and LSFM is highly promising on paper, the technical complexity has so far remained experimentally intractable: structured illumination, in its original implementation introduced by Heintzmann^24^ and Gustafsson^23^, requires sequential illumination with a sinusoidal interference pattern with different phases and orientations (**Figure 1A**). For Light-sheet fluorescence microscopy, typically separate illumination and detection objectives are employed (**Figure 1B**). While a structured light-sheet can be launched through the illumination objective, as has been done in Lattice Light-Sheet Microscopy^25^, the resulting interference pattern cannot be rotated. As a result, the resolution gain is anisotropic (as shown in **Figure 1A**). To illuminate the sample with three differently oriented light-sheets requires multiple illumination objectives^26^. Due to steric hindrance, very little space is left for sample mounting, and glass coverslips would collide with one or more beam paths (**Extended Figure 1**).

**Figure 1.**
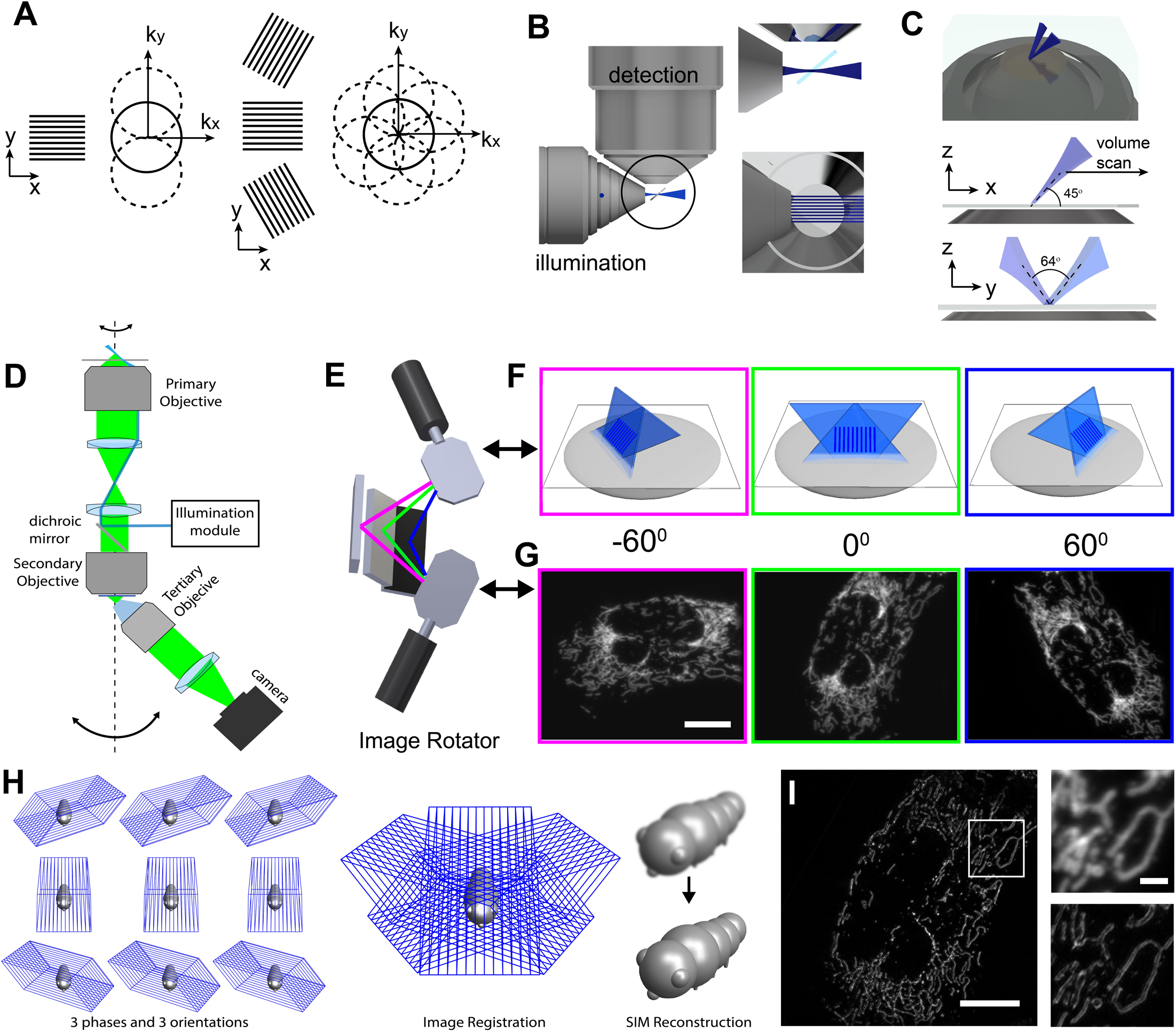
Combination of structured illumination and light-sheet microscopy. **A** Working principle of structured illumination microscopy (SIM): Spatial Fourier components observed by a conventional microscope lie within the solid line circle (i.e., within the passband of the microscope). Using one-dimensional structured illumination (black stripes), additional sample information can be encoded via frequency mixing (dotted circles). Using three differently oriented illumination patterns allows near-isotropic resolution improvement. **B** Geometry of a light-sheet microscope with numerical apertures of 0.7 and 1.1 for illumination and detection. The illumination objective can generate a structured light-sheet, but rotation of illumination pattern as shown in **A** is not possible. **C** Oblique plane microscopy (OPM) using a structured light-sheet. Two mutually coherent light-sheets form an interference pattern along a plane tilted to the coverslip. Volumetric sample information is acquired by scanning the light-sheet laterally. **D** Simplified optical train of an OPM: fluorescence light (green) from the oblique light-sheet (blue) is imaged onto a camera via remote focusing using a secondary and tilted tertiary objective. Arrows indicate rotation of the light-sheet and the detection path. **E** Schematic depiction of the image rotator, which is used in lieu of mechanical rotation of the light-sheet and detection optics. Depending on optical path selection (magenta, green and blue lines), an image rotation of -60,0 and 60 degrees around the optical axis results. **F** Schematic depiction of azimuthal rotation of the structured oblique light-sheets by the image rotator. **G** Maximum intensity projections of an U2OS cell labeled for GFP-OMP25 imaged volumetrically under three different orientations of the structured oblique light-sheet. **H** For Oblique plane structured illumination microscopy (OPSIM), three stacks under different phases of the structured light-sheet are acquired for each azimuthal orientation. The different orientations are computationally registered to each other, and a SIM reconstruction is performed. **I** Mitochondria labeled with GFP-OMP25 in an U2OS cell as imaged by OPSIM. The insets show enlarged versions of the boxed area in **I**, as imaged by OPM (above) and OPSIM (below). Scale Bars: **G** and **I**: 10 microns; inset in **I**: 2 microns

Somewhat surprisingly, there is a solution using less objectives that allows the unconstrained combination of SIM and LSFM, namely by using Oblique Plane Microscopy (OPM)^27^ as the light-sheet platform. Briefly, in OPM, the specimen is illuminated with an obliquely launched light-sheet, and the fluorescence is captured by the same objective. Here, to combine OPM with SIM, we launch two mutually coherent light-sheets at an incidence angle of 45 degrees from a high numerical aperture (NA=1.35) objective (**Figure 1C**). By overlapping their beam waists, a light-sheet with a one-dimensional interference pattern is produced (see also **Extended Figure 2**). To minimize light-losses, the fluorescence light stemming from the tilted structured light-sheet is captured with a recently developed solid immersion tertiary objective (**Figure 1D**)^28^. Unlike the conventional light-sheet geometry (**Figure 1B**), the solid angle available with the primary objective permits rotation of the obliquely launched light-sheet at will (curved arrow around the dotted line in **Figure 1D**), and importantly, there is no steric hindrance or aperture limitation. Practically, however, the detection path would need to co-rotate to stay in alignment with the oblique plane spanned by the structured light-sheet. Conceptually, the whole optical train, or the sample, could be mechanically rotated, albeit at the price of increased technological complexity and slow acquisition speed.

As a high-speed and experimentally tractable alternative to mechanical rotation, we introduce an image rotator, which we designed to perform discrete rotation steps in millisecond time intervals. In contrast to a previously developed image flipper^29^ that is capable of 0-or 180-degree image rotation states, our system can rotate an image under multiple, freely adjustable angles. The system was optimized to work with commercially available galvanometric scanning mirrors (**Figure 1E, Supplementary Note 1, Extended Figure 3**). The image rotator fits in an image space of a high resolution OPM system without requiring any additional relay lens systems or changes to the employed scan and tube lenses (**Methods** and see also **Extended Figure 4** for a schematic drawing of the full setup). In operation, our image rotator can adjust the structured light-sheets to azimuthal orientations of -60, 0, and 60 degrees (**Figure 1F**) as required for an isotropic lateral resolution enhancement by SIM. At the same time, the returning fluorescence light is “de-rotated” by the image rotator such that it is always in alignment with the remote focus system (e.g., to the angled tertiary objective, see also **Figure 1D**). **Figure 1G** shows maximum intensity projections of three volumes of a U2OS cell, transfected with a marker of the outer mitochondrial membrane, GFP-OMP25, imaged under three azimuthal orientations of the structured light-sheet as shown in **Figure 1F**.

To perform oblique plane structured illumination microscopy, or OPSIM for short, we acquire three stacks under different phases of the illumination pattern for each azimuthal orientation (**Figure 1H**). Unlike conventional SIM, the image stacks between the three orientations are rotated to each other (see also **Figure 1G**). This requires computational registration of the three views to each other (**Methods and Extended Figure 5-6**). For the work presented here, we then perform a 2D SIM reconstruction for each X-Y layer on the registered stacks. The SIM reconstruction then effectively fuses the three registered views (**Methods**). **Figure 1I** shows an OPSIM reconstruction of the same U2OS cell as shown in **Figure 1F**. The insets show that in OPSIM, the outer membranes are resolved, making the mitochondria appear hollow, a feature that is not resolved with conventional OPM.

To evaluate the resolution of OPM and compare its image performance, we imaged beads, analyzed their point spread functions (PSF), performed comparative imaging of small biological structures, and employed image decorrelation analysis on a biological sample. Starting with 100nm fluorescent beads in the normal OPM mode (**Figure 2A**), we measured a lateral resolution of 297± 20nm in y and 338± 15nm in the x-direction, respectively. Such anisotropy is typical in OPM due to light-losses induced by the tilting of the tertiary objective. Importantly, the resolution extension by SIM acts in the y-direction, hence it doubles the OPM resolution in the direction it is already highest. For OPSIM, we obtain a resolution of 138 ± 12nm and 148 ± 10nm (average over the full field of view shown in **Supplementary Figure 1A-B**, ± standard deviation) for x and y resolution, respectively (**Figure 2B**). Thus, multidirectional structured illumination not only improves the resolution, but also rectifies the anisotropy of the OPM PSF. In **Figures 2C-F** we compare the 3D PSF of OPSIM to Lattice Light-sheet microscopy (LLSM) with one-directional structured illumination, a Nikon SoRA spinning disk, and a Visitech instant SIM system. Notably, the LLSM PSF shows a high anisotropy in the lateral direction, as only one SIM pattern orientation is possible. However, the axial resolution is the highest of the four PSFs. The OPSIM PSF is similar in its shape and dimensions to the one of the SoRA and iSIM instruments. A comparison of the Full width half maxima for each PSF is provided in **Supplementary Table 1**.

**Figure 2.**
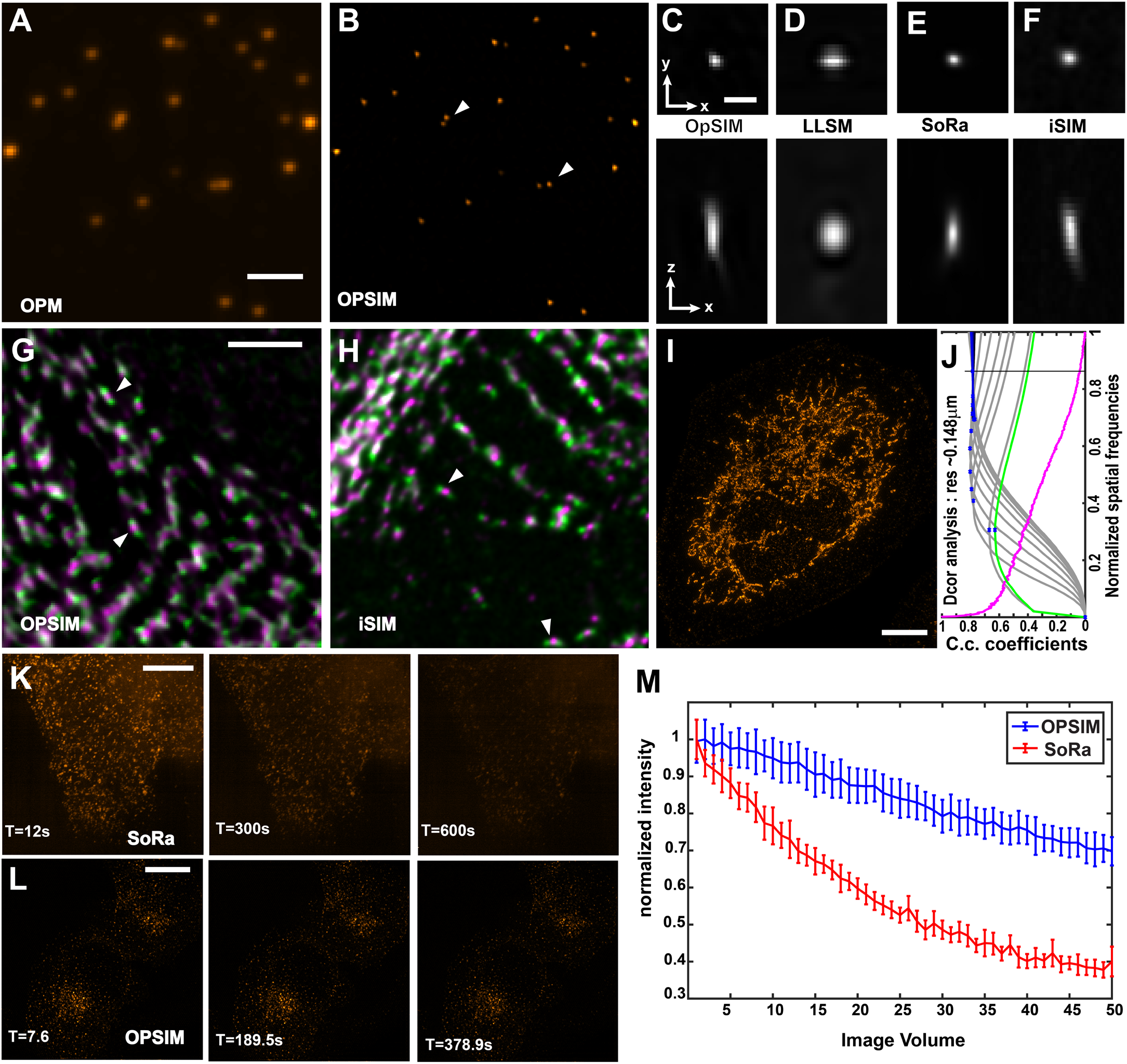
Resolution and performance of oblique plane structured illumination microscopy. **A-B** 100nm fluorescent microspheres as imaged with oblique plane microscopy (OPM) and with oblique plane structured illumination microscopy (OPSIM), respectively. White triangles point at two closely spaced beads that are only resolved with OPSIM. **C-F** Point spread functions for OPSIM, lattice light-sheet microscopy with one-directional structured illumination, a SoRa spinning disk, and an instant structured illumination microscope (iSIM). **G&H** Myosin IIA-GFP motors (green) and Myosin IIA rods (magenta, Alexa 561-conjugated antibody) in a cardiomyocyte as imaged with OPSIM and iSIM. White triangles point at Myosin motor pairs with an interdigitated rod domain that are resolved by each imaging modality. **I** A U2OS cell immunofluorescently labeled for MIC60 with Alexa 488 and imaged with OPSIM (maximum intensity projection). **J** Decorrelation analysis of the cell shown in **I**. Green line: decorrelation function before high-pass filtering, gray lines: decorrelation functions with high-pass filtering, blue dots: local maxima, magenta line: radial average of power spectrum of **I**, C.c.: cross-correlation. **K** Clathrin coated vesicles, labeled for AP2-eGFP, volumetrically imaged over 50 timepoints with a SoRa spinning disk. **L** Clathrin coated vesicles, labeled with AP2-eGFP, imaged over 50 timepoints using OPSIM. **M** Normalized fluorescence intensities of the 20 brightest vesicles over time for OPSIM and SoRa imaging. Scale bars: **A**,**G** 2 microns; **C** 500nm; **I**,**K**,**L** 10 microns.

We next sought to demonstrate the enhanced resolution on biological targets. In iPSC-derived cardiomyocytes, Myosin IIA forms filaments wherein individual Myosin IIA motor domains are spaced ∼300nm apart, separated by a rod domain in between^30^. When separately labeled, Myosin IIA motors and rods are only visible with super-resolution fluorescence microscopy^30^. In this scenario, the motors appear as two puncta flanking one another, and the rod appears as a single punctum. These three resulting puncta are roughly 150nm-spaced, representing a super-resolution “ruler” for SIM microscopy. **Figures 2G-H** show cardiomyocytes transfected with Myosin IIA-GFP (whereby GFP is attached to the Myosin IIA motor) and stained for Myosin IIA rod domains with Alexa Fluor 561 conjugated antibodies. As shown, both OPSIM and instant SIM are capable of successfully resolving the Myosin IIA motors and the corresponding interdigitated rod domain. For a more quantitative measurement of resolution within a cell, we also performed image decorrelation analysis^31^ on a U2OS cell (**Figure 2I-J**), fixed and labeled with an Alexa Fluor 488 conjugated antibody targeting MIC60, a mitochondrial inner membrane protein known to mark sites of cristae invagination^32^. The image decorrelation resolution estimate of 148nm is in good agreement with the FWHM measurements of fluorescent nanospheres.

One promise of combining LSFM and SIM is to reduce the rate of photobleaching. To investigate this, we imaged eGFP-tagged AP2 volumetrically in parental human retinal pigmented epithelium (ARPE-19) with OPSIM and a SoRa spinning disk over 50 timepoints, while keeping the total stack acquisition time and volumetric coverage close to identical (see also **Supplementary Table 2** for acquisition parameters). **Figures 2K-L** show the first, 25^th^ and 50^th^ timepoint. As can be seen from those timeframes, and from **Figure 2M** where the average intensity of the twenty brightest spots is compared, OPSIM offers a notably lower rate of photo-bleaching.

To demonstrate the potential of OPSIM for biological discovery, we first imaged the cytoskeleton in hTERT immortalized RPE cells and cardiomyocytes. **Figures 3A-B** show the actin cytoskeleton and vimentin intermediate network in hTERT RPE cells, respectively, as imaged with OPSIM. A comparison to conventional OPM shows that finer details of the filamentous networks become apparent, such as twists and branching points. **Figure 3C-D** shows a cardiomyocyte labeled with phalloidin-488 (which labels actin filaments) and Alexa Fluor 561 conjugated antibodies targeting alpha-actinin 2 (which labels the sarcomeric Z-lines). OPSIM imaging shows an enrichment of phalloidin in the z-disks that colocalizes with the actinin labeling (see also **extended Figure 7**). We also evaluated denser samples, which are typically more challenging to image with SIM due to reconstruction artifacts owing to out-of-focus blur. Here we imaged fixed, 20-micron thick slices of a mouse spinal cord using Alexa Fluor 488 and 555 conjugated antibodies, respectively, targeting neurofilaments and proteolipid protein (PLP), which labels the myelin sheath (**Figure 3E**). OPSIM revealed puncta of the PLP labeling, which is expected due to anchoring of PLP in between the myelin sheath membrane^33^. Notably, a standard confocal image does not resolve this structure of the myelin sheath clearly, whereas the SoRa spinning disk also revealed a similar distribution (**Extended Figures 8-9**).

**Figure 3.**
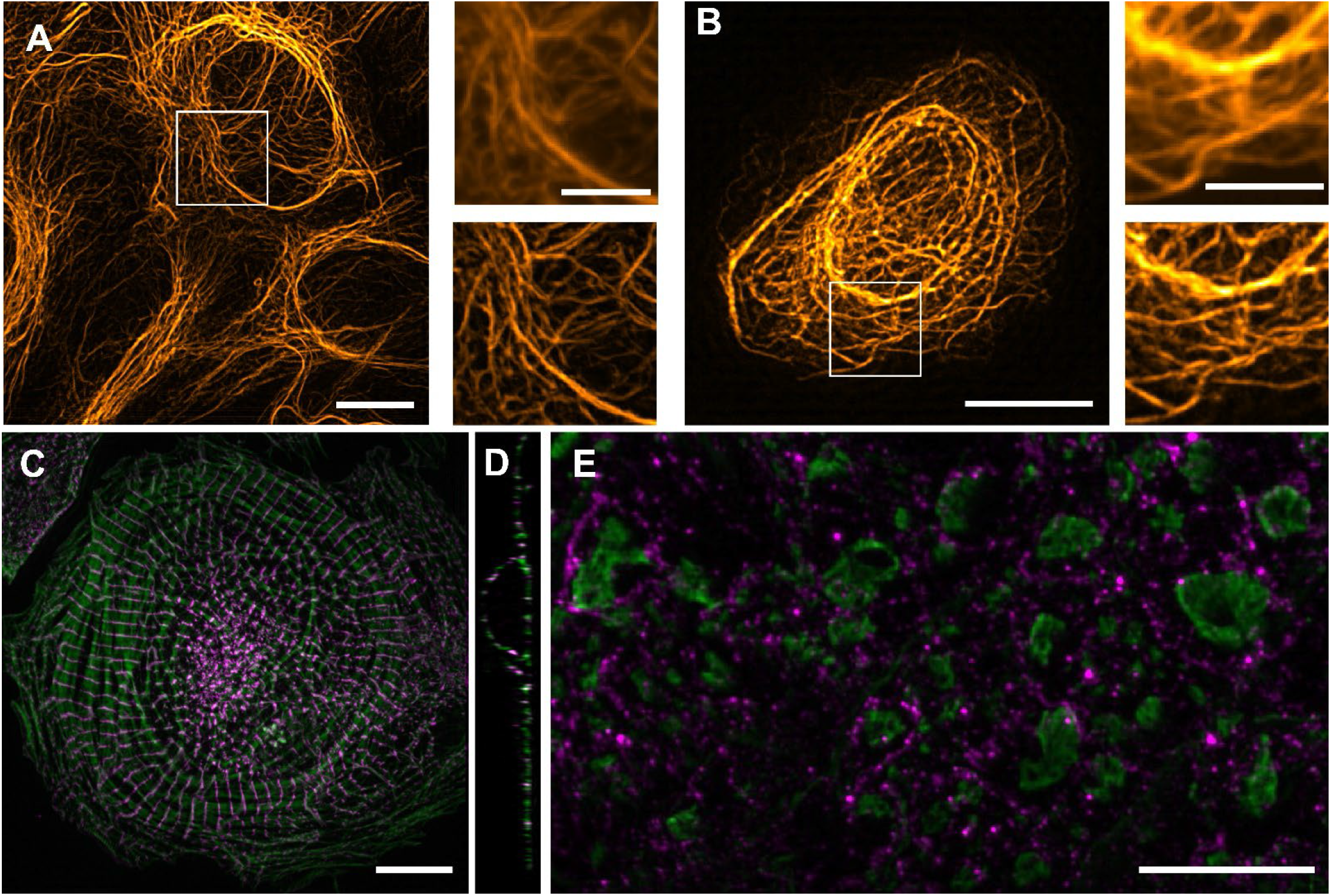
Imaging of cellular samples with OPSIM. **A** The actin cytoskeleton in an hTERT RPE cell as imaged by OPMSIM. Insets show the boxed region in **A**, as imaged by OPM (above) and OPSIM (below). **B** Vimentin in an hTERT RPE-1 cell as imaged by OPMSIM. The insets show enlarged versions of the boxed area in **B**, as imaged by OPM (above) and OPSIM (below). **C** Cardiomyocyte labeled for actin (green, Alexa Fluor 488-phalloidin) and alpha-actinin 2 (magenta, Alexa 561-conjugated antibody), as imaged by OPSIM. **D** Cross-sectional view (single slice) of the cardiomyocyte shown in **C. E** Spinal cord slice, immunofluorescently labeled for neurofilament (green) and myelin (magenta), as imaged by OPSIM. **A**,**B**,**C**,**E** show maximum intensity projections. Scale bars: **A**,**B**,**C**,**E**: 10 microns; insets of **A** and **B**: 5 microns

To illustrate the potential for rapid volumetric imaging, we captured the dynamics of clathrin coated vesicles and mitochondria. **Figures 4A-B** show a maximum intensity projection of an ARPE-19 cell, labeled with AP2-eGFP, as imaged by OPSIM at 0.86 Hz volumetric rate over 40 timepoints. **Figures 4C-D** show a montage of four selected time points that highlight vesicle dynamics (see also **Movie 1**). Notably, OPSIM enables observation of vesicle movements in 3D (**Figure 4D, Movie 1**) at a temporal resolution typically employed for clathrin mediated endocytosis studies using TIRF microscopy^34, 35^. However, in contrast to TIRF, the vesicles can now be studied across the entirety of the cell, and axial movements will not lead to a loss of the vesicle, as is the case when a vesicle leaves the evanescent field in TIRF microscopy^36^. **Figure 4E** shows an U2OS cell, labeled with GFP-OMP25, that was imaged with OPSIM at a 1.2 Hz volumetric acquisition rate over 38 timepoints. **Figures 4F-G** show a montage of four selected time points for two areas, which highlight rapid protrusion (**Figure 4F**) and retraction (**Figure 4G**) events of mitochondria. OPSIM resolves these dynamics without apparent motion blurring (**Movie 2**). The volumetric rate in OPSIM can be further improved by shrinking the observation volume (see also **Movie 3**, which was acquired under a 1.4Hz volume rate).

**Figure 4.**
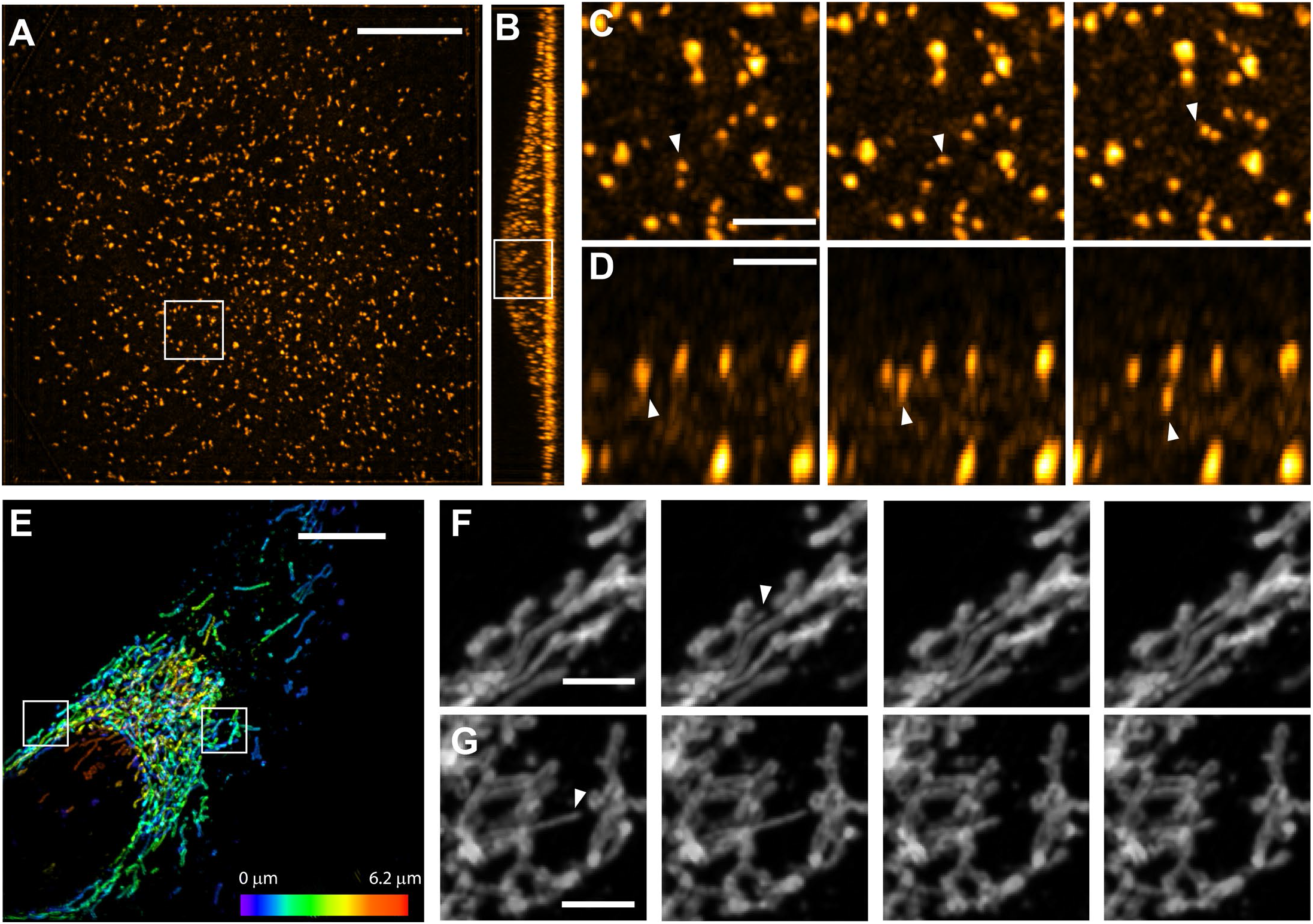
Dynamic volumetric imaging with OPSIM. **A** An ARPE-19 cell, labeled for AP2-eGFP, as imaged volumetrically by OPSIM at 0.86Hz. A maximum intensity projection is shown. **B** cross sectional view of the ARPE cell in **A** (maximum intensity projection). **C** Three selected timepoints of the boxed area in **A**. Temporal separation between each slice is 3.5s **D** Three selected timepoints of the boxed area in **B**. Temporal separation between each slice is 2.3s White triangles point at movement of a clathrin coated vesicle. **E** Mitochondria in an U2OS cell, as imaged by OPSIM at a volumetric rate of 1.2 Hz. Maximum intensity projection with height above the coverslip encoded in color. **F** Selected timepoints of the boxed region on the left in **E**. Temporal separation between each slice is 0.82s. White triangle marks the tip of a protruding mitochondrion. **G** Selected timepoints of the boxed region on the right in **E**. Temporal separation between each slice is 1.64s White triangle marks the tip of a retracting mitochondrion. Scale Bars: **A**,**E**: 10 microns. **C-D, F-G**: 2 microns.

In summary, we show to our knowledge the first practical implementation of LSFM with multi-directional structured illumination. In striking contrast to previous efforts, our solution is compact and experimentally tractable: only two new modules, each of which integrate into existing OPM platforms without requiring additional optics, are needed. As SIM requires high optical precision and raw data that is diffraction limited, it was not known if an image rotator would introduce pointing errors that would offset the delicate alignment of a high resolution OPM system, jeopardize the positioning of the excitation beams at the edge of the objectives’ pupil, or distort the three differently oriented volumes. However, the fact that all three orientations feature diffraction limited point-spread functions, and the corresponding volumes can be registered with linear affine transformations to the precision of a SIM reconstruction (i.e., point sources overlap tightly across the volume at the doubled resolution) indicates that the OPSIM described herein achieves sufficient optical precision. Furthermore, it was not known if the extensive pre-processing of the data (i.e., shearing, rotating, and registering the data, all of which involve interpolation operations) could interfere with a SIM reconstruction. The Fourier transforms of pre-processed x-y slices feature sharp peaks for the Fourier components of the illumination pattern (**Supplementary Figure 2**), and the SIM reconstructions are void of artifacts beyond what is typically occurring in a conventional SIM system (**Supplementary Figure 1**), also in regard to motion blurring. Overall, the quality of the reconstructions indicates that the OPSIM optical system and corresponding data preprocessing are precise enough to perform faithful SIM reconstructions.

More could be done to improve the z-resolution using optical (e.g., 3D SIM pattern, axially swept light-sheet microscopy^37, 38^), spectroscopic (e.g., reversibly photoswitching fluorophores), and computational (e.g., performing the SIM reconstruction in 3D, 3D image fusion, iterative 3D deconvolution, convolutional neuronal networks for image restoration) methods. In our current implementation, we did not pursue any of these options as we wanted to establish a first proof of concept for OPSIM and its basic operation principles. The use of 2D SIM, i.e., illumination with a one-dimensional pattern and slice by slice processing, simplified the illumination module, and made the image acquisition shorter and robust.

As implemented, OPSIM does not reach the resolution of a 3D SIM system with sinusoidal illumination patterns. This is in part due to the reduced resolution in an OPM detection path compared to pure widefield detection utilized in a 3D SIM system, and the lower illumination NA that can be realized for an oblique structured light-sheet (see also **Extended Figure 10**). However, the resolution of OPSIM is comparable to SIM techniques known as Image Scanning Microscopy or instant SIM. What sets OPSIM apart from previous SIM systems is its light-sheet excitation, which avoids unnecessarily exciting portions of the sample above and below the focal plane. As such, lower photo-bleaching rates are expected, which we confirmed experimentally. Additionally, the physical optical sectioning of OPSIM, compared to the widefield detection of traditional 3D SIM systems, is expected to make it more robust against reconstruction artifacts when imaging thick samples.

OPSIM has the potential to accelerate volumetric SIM imaging, owing to the low inertia volumetric scanning mechanism^39^. Indeed, the presented volumetric imaging at ∼1.4 Hz is among the fastest volumetric SIM rates reported to our knowledge. In contrast, conventional SIM systems are more fundamentally limited by mechanical sample scanning, and photo-bleaching due to the lack of excitation confinement. OPSIM, with the advantages of light-sheet excitation and its low inertia scan mechanism, may open new opportunities to accelerate volumetric SIM imaging. Even faster volumetric imaging than demonstrated in this manuscript might be desirable for many dynamic, three-dimensional processes. Indeed, even at 1.4 Hz acquisition rate, rapidly moving clathrin coated vesicles exhibited motion artifacts, in contrast to stationary vesicles that appear as round spots (See also **Movie 3**). OPSIM has the potential to be operated even faster by: using faster cameras, employing only two image rotations (0 and 90) at the expense of resolution isotropy and by combining OPSIM with the recently invented projection imaging technique^40^. The latter approach, i.e., projecting volumes under SIM illumination into a common reference frame followed by a 2D SIM reconstruction, could speed up acquisition times by up to two orders of magnitude. However, only projected views of a 3D scenery can be obtained this way.

In conclusion, the combination of LSFM and SIM holds promise for volumetric imaging with high spatiotemporal resolution and low levels of photo-toxicity. The involved technical complexity has so far rendered such systems overly complex and ill-suited for common sample mounting methods. With OPSIM, we have introduced a practical solution that combines rapid volumetric imaging at 150nm lateral resolution with the advantages of light-sheet excitation and the ease of conventional sample mounting. As such, we think OPSIM will extend the number of biological questions that can be studied with volumetric fluorescence microscopy.

## Acknowledgments

We would like to thank the National Institutes of Health (1R01DK127589, 1R21HD105189, 5P30CA142543, and U54CA268072 to K.M.D. MIRA to D.T.B. R35 GM125028, R35 GM133522 to R.F., R35GM137894 to JRF.). American heart Association Graduate Student Fellowship to J.B.H. (AHA 836090). P.R. received funding from the Investissements d’Avenir French Government program managed by the French National Research Agency (ANR-16-CONV-0001) and from Excellence Initiative of Aix-Marseille University -A*MIDEX.”

## Author Contributions

R.F. conceived the idea of OPSIM and the image rotator. B.C. mathematically described the image rotator and B.J.C performed numerical simulations and practical experiments. B.C., B.J.C. and R.F. designed an experimentally tractable image rotator. B.C build the image rotator. R.F. build the microscope. R.F. and B.J.C. acquired experimental data. R.F., P.B., B.J.C. and B.C. performed data processing. P.R. and F.Z. wrote the fine registration algorithm. R.F., B.J.C, P.B and D.S. wrote the SIM reconstruction software. M.M., J.H., J.F., E.S., K.D.M., D.B., C.W.Z. and C.L.Z provided biological samples. E.S. and J.H. performed imaging with the Sora and iSIM system, respectively.

## Data Availability

All data presented herein are available from the corresponding author upon request. Example OPSIM reconstructions are provided under: https://zenodo.org/record/6481084#.YmVM-7lOmHs

## Code Availability

The MATLAB scripts used in this manuscript are available under: https://github.com/AdvancedImagingUTSW/manuscripts/tree/main/2022-chen Example OPSIM raw data is provided under: https://zenodo.org/record/6481084#.YmVM-7lOmHs The Python scripts are available under: https://github.com/QI2lab/mcSIM. The version of the code used here is archived on Zenodo (https://doi.org/10.5281/zenodo.6419901).

## Methods

### Oblique Plane Structured Illumination Microscope

Compared to our previously published OPM system^28^, OPSIM employs a 45 degrees tilt of the light-sheet and the tertiary imaging arm and incorporates two new modules, an image rotator, and a Michelson interferometer to generate and rotate the structured light-sheet. The OPSIM optical train (see also **Extended Figure 4** for a schematic drawing) uses a Nikon 1.35/100X silicone immersion objective lens, followed by a 200mm tube lens (ITL200, Thorlabs), two scan lenses (CLS-SL, Thorlabs) bracketed around a galvanometric mirror (6215, Cambridge Technologies), a custom tube lens with an effective focal length of 357mm (consisting of an AC508-750-A and an AC508-500-A achromatic lens, Thorlabs), a Nikon 40X/0.95 air objective, a solid state tertiary objective (AMS-AGY objective - v1.0, Applied Scientific Instrumentation), a 200mm tube lens (ITL 200, Thorlabs) and a sCMOS camera (Flash 4.0, Hamamatsu). A motorized flipper mount (MFF101, Thorlabs) was used to switch emission filters (FF01-525/50-25 and FF01-593/LP-25, Semrock) in front of the camera for sequential dual color imaging.

The microscope uses a 488nm and 561nm diode pumped solid state laser (Sapphire 488-300 and OBIS 561-150LS, Coherent Inc) for illumination. The intensity of the 561nm laser is modulated via software control. The 488nm laser is intensity modulated with an external acousto-optical filter (AOM-402AF1, IntraAction). After combining both laser lines with a beam splitter, a mechanical shutter (SH05, Thorlabs) is used to turn the laser illumination on and off. The laser beam was cleaned up with a pinhole (30-micron diameter, Thorlabs) in between a telescope (AC254-50A and AC254-150A, Thorlabs), followed by a beam expander (GBE03A, Thorlabs). S-polarization of the laser beams was adjusted with a half waveplate before entering the Michelson interferometer. Of note, the polarization was further rotated by the image rotator, ensuring S-polarization for each illumination direction. The expanded laser beam then entered the Michelson interferometer described below.

The data acquisition computer was a Colfax International ProEdge SXT9800 Workstation. The control software was developed using a 64-bit version of LabView 2016 equipped with the LabView Run-Time Engine, Vision Development Module and Vision Run-Time Module (National Instruments). The Software communicated with the camera via the DCAM-API for the Active Silicon Firebird frame-grabber and delivered TTL triggers and analog voltage signals through a field programmable gate array (PCIe 7852R, National Instruments). These triggers and analog voltage signals control the optical shutter, galvanometer mirror scanning, steering of the image rotator galvos, phase stepping of the piezo in the Michelson interferometer, and camera fire and external trigger. The control software can be requested from the corresponding authors and will be distributed under an MTA with the University of Texas Southwestern Medical Center.

### Michelson Interferometer

A cylindrical lens (ACY254-100A, Thorlabs) is used to focus a line onto a pair of mirrors, conjugated to the sample plane via a nonpolarizing beam splitter. Both mirrors can be mechanically controlled in tip and tilt, which allows to adjust the incidence and azimuthal angle of each light-sheet individually. One of the mirrors is mounted on a high-speed piezo actuator (Nano-OP30HS, Mad City labs) for phase stepping. An adjustable slit (VA100, Thorlabs), placed at one focal length in front of the cylindrical lens, is used to control the divergence of the light-sheets.

To couple the structured light-sheet into the OPSIM system, the laser light passes through an achromatic doublet lens (AC254-100A), is reflected by a motorized (PIA25, Thorlabs) tip tilt mirror, and reflected into the OPM train by a dichroic mirror (Di03-R405/488/561/635-t3, Semrock).

The motorized mirror is conjugate to the pupil plane of the primary objective. That way, the motorized mirror can be used to shift the light-sheet laterally and align it with the oblique plane that is in focus with the tertiary imaging system.

### Image Rotator

The image rotator consists of two major components: a pair of galvanometric mirrors (6220H, Cambridge Technologies) and a set of static mirrors (#46-721, Edmund Optics). The galvanometric mirrors were held by custom aluminum mounts and static mirrors by mounts and optical post assemblies from Thorlabs (**Extended Fig. 3**). The Galvos were placed with the angle between the axis of 60°, and a separation of 76mm between them. For mechanical rotation angles of ± 10° and 0° for the two Galvos, the laser beam or fluorescence image will be reflected off the first Galvo onto one of the static mirrors, then reflected to the second Galvo. De-scanned by the second Galvo, the optical propagation axis of the output will remain stationary, resulting in the laser beam and fluorescence image being rotated by ± 60° and 0°. Two additional folding mirrors under the galvanometric mirrors were used to obtain collinearity of input and output. We used 3D Mirror Transformation Matrices and Zemax to analyze and present the working principle of the image rotator and calculate the discrete rotation angles (**Supplementary Note 1**). To find the correct position of the Galvo and static mirrors, we split an alignment laser into two beams and injected them from opposing sides into the image rotator. Then we adjusted the Galvo mirrors to make the two laser beams overlap at each rotation status. The stationary mirrors were placed such that they coincide with the beam overlap. Correct alignment was further confirmed by a series of irises placed on either output of the image rotator. This allowed us to minimize pointing and translation errors by carefully aligning the tip and tilt of the stationary mirrors. This alignment was performed outside of the OPM microscope on a large enough optical bench to allow ∼1m of free space on either side of the rotator. The aligned image rotator was then integrated into the image space between the first tube and scan lens (see also **Extended Figure 4**).

The field of view in OPSIM is mainly limited by the image rotator and is governed by the size of the galvo mirrors and the maximum inclination of the Galvo mirror compared to the incident light. Further, the divergence angle of the light in the image space also plays a factor in limiting the field of view. In our system, all these factors combined limited the field of view to about 60×60 microns. The field of view could be increased by using larger galvo mirrors, and by making the maximum inclination of the Galvo mirror smaller.

### Lattice Light-Sheet Microscopy

The Lattice Light-Sheet SIM imaging was performed by a home-built instrument that was setup Harvard medical School following the design outlined within the seminal LLSM publication^25^.

### Instant SIM imaging

The instant SIM (iSIM) imaging was performed with a Visitech iSIM using a Nikon SR HP Apo TIRF 100x oil immersion objective (model number MRD01997) at 1.5X zoom with NA=1.49. The laser lines used for imaging excite at 488-491nm (green) and 561nm (red). Images were captured using a Hamamatsu ORCA-Fusion Digital CMOS camera (model C14440-20UP) with a 0.1um axial step size. Images were deconvolved using Microvolution software (Cupertino, CA) installed in FIJI (Fiji Is Just ImageJ) over 20 iterations.

### SoRa Spinning disk imaging

SoRa spinning disk imaging was performed on a Nikon CSU-W1 microscope equipped with an 100x objective of NA 1.45. Deconvolution in the NIS-Elements AR 5.21.03 proprietary software was performed over 10 iterations.

### Confocal Microscopy

Confocal microscopy was performed on a Nikon A1R microscope equipped with an 60x objective of NA 1.4.

### Mammalian Cell Culture

U2OS cells were cultured in DMEM supplemented with 10% Fetal Bovine Serum FBS (Sigma), 25mM HEPES, 100units/mL penicillin, and 100ug/mL streptomycin. GFP-OMP25 (Addgene #141150, a kind of Gia Voeltz^41^) was transiently transfected using Lipofectamine 3000 (ThermoFisher) according to manufacturers directions and cells were passaged to CellVis glass bottom microscope dishes. Cells were grown to ∼50% confluency in glass bottom dishes and fixed in 4% paraformaldehyde solution in PBS (15 minutes, room temperature). Fixed cells were permeabilized (0.1% Triton X-100 in PBS), blocked (10% FBS and 0.1% Triton X-100 in PBS), and then immunolabeled with anti-MIC60 antibody (ProteinTech 10179-1-AP) and donkey anti-rabbit-AlexaFluor488 (ThermoFisher A-21206).

Cardiomyocytes used in this study are human induced pluripotent stem cell-derived cardiomyocytes purchased directly from Cellular Dynamics International (iCell cardiomyocytes, Madison, WI, https://www.fujifilmcdi.com/icell-cardiomyocytes-2-11713). Cells were cultured as per manufacturers instructions - briefly, upon thawing, cardiomyocyte maintenance medium is added to cardiac myocytes every 48 hours. Cardiomyocyte transfections were performed with Viafect (promega) using a protocol whereby a viafect:plasmid mixture is added to cells at a 6:1 uL viafect:ug plasmid ratio for 24 hours^42^. Cardiomyocytes were plated on fibronectin-coated, Cell Vis glass-bottom dishes for imaging. Cardiomyocyte staining was performed following fixation and permeabilization similar to U2OS cells above. Phalloidin staining (if performed) was performed prior to blocking by incubating cells with 15uL phalloidin:185uL 1X PBS for 2 hours at room temperature. Cardiomyocytes were blocked in buffer containing 10% bovine serum albumin (BSA) in PBS. Primary and secondary antibodies were added to cells in blocking buffer for 1 hour, 45 minutes and 1 hour, respectively, at room temperature. For the phalloidin-actinin stain, alpha-actinin-2 was labeled using the anti-sarcomeric primary antibody EA-53 (Abcam, mouse) and Alexa Fluor 561 secondary (goat anti-mouse). For the NMMIIA stain, the NMMIIA rods were labeled using 909801 anti-non-muscle-myosin IIA heavy chain primary antibody (Biolegend, rabbit) and Alexa 561 secondary (goat anti-rabbit).

Parental human retinal pigmented epithelium (ARPE-19) cells obtained from ATCC were infected to stably express eGFP-labeled AP2 as previoduly described^34^. Separately, hTERT immortalized human retinal pigment cells (hTERT RPE-1) obtained from ATCC were modified to express fluorescent protein tagged isoforms of vimentin and microtubules at their native genomic loci. Both retinal epithelial cell lines were cultured in DMEM/F12 supplemented with 10% FBS at 37°C in a 5% CO2 atmosphere and imaged on 35 mm glass bottom dishes (MedTek Corp.)

### Spinal cord slice samples preparation

The spinal cord was embedded in OCT cryostat-embedding compound (Tissue-Tek, Torrance, CA), cut into 20-μm thick sections on a cryostat (Leica Microsystems, Germany) at −23°C, and mounted on gelatinized slides (Thermo Fisher). Primary antibodies were prepared from rabbit anti-NF200 (neurofilament marker; Sigma) at 1:500, and mouse monoclonal primary antibody against myelin basic protein antibody (Millipore) at 1:200 dilution. Secondary antibodies were goat anti-rabbit and anti-mouse Alexa 488 and Alexa 555 conjugated fluorescence at 1:500 (Millipore). The spinal cord sections were directly coverslipped with Dapi mounting media (Vector, Burlingame, CA) and analyzed on the microscope right afterwards.

### Data Pre-Processing

A Matlab program performs the pre-processing steps of the raw data prior to a SIM reconstruction (see also **Extended Figure 5-6** for a graphical representation of the steps): Each data volume is de-skewed and numerically rotated by 45 degrees into the coverslip coordinate system (z being along the optical axis of the primary objective, and normal to the coverslip). The datasets for each illumination direction are then rotated to each other along the z-axis and the volumes are shifted onto each other by using cross-correlations of maximum intensity projections (top and side views). This roughly aligns the data volumes from each viewing direction to each other on a pixel level basis. To achieve a more precise registration accounting additionally for 3D rotation and skew, sufficiently accurate at the higher SIM resolution, pairwise multiscale affine registration was used. Data volumes were downsampled progresssively in pyramidal fashion at 1x, 2x, 4x, 8x and 16x. The 4×4 affine transformation matrix was then learnt first by aligning data volumes at the coarsest 16x scale. The fitted parameters then initialises the affine matrix for the finer 8x level and so on to the 1x scale. To achieve robustness to potential intensity and contrast variation, data volumes were regarded as ‘multimodal’ and Matte’s mutual information metric based on comparing the pixel intensity histogram is used instead of the raw pixel intensity differences. To register the three different directions, direction 0 and direction 2 were registered as described to direction 1 (see also **Extended Figure 6** for naming convention).

### Structured illumination reconstruction

SIM processing was performed slice-by-slice (i.e., one x-y plane, parallel to the coverslip, at the time) using the pre-processed data. The algorithm follows the principles outlined by Heintzmann and Gustafsson, in the sense that for each illumination direction, a linear equation system is solved in Fourier space on a pixel-by-pixel basis to separate the different information bands (i.e. a zero order band, corresponding to a conventional OPM spectrum, and two sidebands containing higher frequency Fourier components). The sidebands are then numerically shifted to their corresponding position in reciprocal space. After combing the different information bands, averaging them in regions of overlap and filtering them with the respective transfer function (i.e., Wiener deconvolution obtained from an experimental PSF), an apodization function is applied before an inverse Fourier transform yields the SIM reconstruction in real space.

The Heintzmann/Gustaffson style SIM processing code is implemented in python and available on GitHub^43^ It includes several options for performing the SIM reconstruction, largely following either OpenSIM^44^ or fairSIM^45^. Various functions are provided for estimating the SIM parameters. For example, the pattern frequencies can be identified directly from Fourier space peaks or using the cross-correlation of separated bands. Pattern phases are identified using either the cross-correlation with a sinusoidal pattern or the iterative method of Wicker^46^. The modulation depth can be estimated by comparing the overlap region of the separated bands or by fitting the sample power spectral density. The code also contains several options for achieving optical sectioning, including implementing the method of Wilson et al^47^ and combining optical sectioning with super resolution reconstruction by filling of the missing cone, as outlined by O’Halloran and Shaw^48, 49^. The latter option proved helpful for OPSIM, as it reduced background blur. Finally, the code includes several quality checks, such as graphical readout of the modulation peak intensity, the modulation peak fitting, and the modulation contrast to noise ratio^50, 51^.

We implemented a second reconstruction algorithm in Matlab that employs Richardson Lucy (RL) deconvolution^52^ instead of the traditional Wiener deconvolution, over a small number of iterations (2-5) using a synthetic PSF. This was developed to enable a parameter-free reconstruction, as the setting of a Wiener constant and the use of an apodization function is not necessary. Furthermore, the high-quality experimental PSF needed for a Wiener deconvolution can be replaced with a synthetic PSF, which was helpful when exploring other color channels (we had only acquired experimental PSFs for the green channel). Lastly, we found that the RL deconvolution also prevents undershoots, which often plaque traditional SIM reconstructions (see also **Supplementary Figure 1 C-D)**.

The processing was implemented as follows: before unmixing the information bands for one image plane, the 2D raw image data was RL deconvolved over 5 iterations. This served to flatten the spectral response. Each separated information band was truncated to a circular region with a radius of 1/290nm^-1^, which we assumed to be the cut-off frequency (here named Kc) for our OPM system, was shifted to its proper place in Fourier space and added together, with appropriate weighting in the overlap regions. In particular, within a radius of Kc /4, only information from the sidebands was used, which served to fill the missing cone of the zero-order band. The reconstructed, extended object spectrum was inverse Fourier transformed to real space without applying an apodization functions. To compensate ringing and undershoots, two iterations of RL deconvolution were applied to the reconstructed data. Alternatively, we implemented a triangular apodization function, but found in practice that the two iterations of RL deconvolution yielded better results in terms of ringing and undershoot suppression.

Importantly, while RL deconvolution with more iterations (e.g.,10-20) could potentially further improve the resolution in SIM, we tried to avoid this effect here by limiting the number of iterations. Indeed, resolution measurements between a SIM reconstruction using Wiener and RL deconvolution are closely matched: the SIM reconstruction using RL deconvolution yielded 6-9% narrower FWHM in the x and y direction, respectively, compared to the Wiener deconvolution SIM reconstruction. The main difference was the suppression of undershoots (see also **Supplementary Figure 1 C-D**).

While it is tempting to employ more iterations of the RL algorithm, we wanted to avoid this here, so that the reader can clearly delineate how much resolution gain stems from the SIM technique.

