## Supplementary Information for "Resolution doubling in light-sheet microscopy via oblique plane structured illumination"

### Supplementary Notes

#### Supplementary Note 1: Design of the Image Rotator

The conventional ways of rotating an image optically are achieved by mechanically rotating a Dovetail or an Abbe–Koenig prism, or a K-mirror system, which are inserted in the optical path of an imaging system. In such systems, the prism or the whole K-mirror system must be mounted on a mechanical rotation stage, which makes the rotation slow (on the order of 100ms to seconds). Besides, the prisms will introduce aberrations, such as astigmatism, when used with converging light, which can be overcome with a K-mirror assembly. To overcome the speed limitation, the incident beam could conceivably be split into multiple light paths which are then rotated individually. Switching between the different optical paths could be performed by mechanical flip mirrors (slow) or using polarization optics (not suited for unpolarized light). However, such systems are overly complex, and either not fast or not generally applicable.

To address these challenges, we invented a rapid image rotator which can perform a discrete number of rotations whose magnitude can be chosen freely. The core components of the image rotator are a pair of Galvos and a set of static mirrors, one for each desired rotation angle (See also **Figure 1E**). The incident light will be reflected by the first Galvo towards one of the static mirrors, then deflected to the second Galvo. Because of the symmetry of the structure, the collinearity of the output light under different rotation angles is maintained. The number of static mirrors is determined by the rotation angles the application needs. Of note, the rapid image rotator also rotates the polarization of the light, unless the mirrors introduce significant de-polarization. Because of an odd number of mirrors, the overall effect also includes a flip of the image, meaning that there is always a bias (or default) rotation present. Because the rotation speed is only limited by the scanning speed of the Galvo pair, this device can perform image rotations in a matter of milliseconds. **Extended Figure 3** shows a CAD drawing for Image rotator.

To investigate how the different parameters (rotation angle of the galvo, arrangement of the different components) affect the image rotation, we performed analytical calculations. We use 3D Mirror Transformation Matrices and Zemax to analyze and present the working principle of the image rotator and calculate the discrete rotation angles. The angle of the image rotation refers to rotation angle relative to the status when the light travels over the central static mirrors. To simplify the model analysis, we only include the Galvo pair and the static set of mirrors when using the matrix formalism, as shown in **Supplementary Fig. 3**. For an ideal system, we assume both galvo mirrors rotate by the same magnitude $\pm\alpha$. The rotation axis (black dashed line) of Galvo Mirror 1 ($G_1$) is perpendicular to the plane of incident and emergent rays (red arrows). The matrix of $G_1$ can be obtained by rotating a virtual z mirror ($M_z$, surface normal parallel to z axis) around x axis for $(45° - \alpha)$ and then around y axis for $\theta$.

$$\mathrm{G}_1 = \mathrm{R}_\mathrm{y}(\theta)\mathrm{R}_\mathrm{x}(45° - \alpha)\mathrm{M}_z{\mathrm{R}_\mathrm{x}}^T(45° - \alpha){\mathrm{R}_\mathrm{y}}^T(\theta)$$

$$= \begin{bmatrix} \cos^2\theta - \sin 2\alpha \sin^2\theta & \cos 2\alpha \sin\theta & -\sin 2\theta\,(\sin 2\alpha + 1)/2 \\ \cos 2\alpha \sin\theta & \sin 2\alpha & \cos 2\alpha \cos\theta \\ -\sin 2\theta\,(\sin 2\alpha + 1)/2 & \cos 2\alpha \cos\theta & \sin^2\theta - \sin 2\alpha \cos^2\theta \end{bmatrix}$$

where, $\mathrm{M}_\mathrm{z} = \begin{bmatrix} 1 & 0 & 0 \\ 0 & 1 & 0 \\ 0 & 0 & -1 \end{bmatrix}$, $\mathrm{R}_\mathrm{x}(\alpha) = \begin{bmatrix} 1 & 0 & 0 \\ 0 & \cos\alpha & -\sin\alpha \\ 0 & \sin\alpha & \cos\alpha \end{bmatrix}$, $\mathrm{R}_\mathrm{y}(\theta) = \begin{bmatrix} \cos(\theta) & 0 & \sin(\theta) \\ 0 & 1 & 0 \\ -\sin\theta & 0 & \cos(\theta) \end{bmatrix}$.

Similarly, we can have the matrix of Galvo Mirror 2 ($G_2$)

$$\mathrm{G_2} = \mathrm{R_y}(-\theta)\mathrm{R_x}(-45° + \alpha)\mathrm{M_z}\mathrm{R_x}^T(-45° + \alpha)\mathrm{R_y}^T(-\theta)$$

$$= \begin{bmatrix} cos^2\theta - sin\,2\alpha\, sin^2\theta & cos\,2\alpha\, sin\,\theta & sin\,2\theta\,(sin\,2\alpha + 1)/2 \\ cos\,2\alpha\, sin\,\theta & sin\,2\alpha & -cos\,2\alpha\, cos\,\theta \\ sin\,2\theta\,(sin\,2\alpha + 1)/2 & -cos\,2\alpha\, cos\,\theta & sin^2\theta - sin\,2\alpha\, cos^2\theta \end{bmatrix}.$$

According to the vector law of reflection, the relation between the unit vector ($n_1$) of the surface normal of Mirror 1 ($M_1$), incident ray ($k_1$) and reflected ray ($k_2$) is

$$k_2 = k_1 - 2(k_1 \cdot n_1)n_1.$$

$k_1$ and $k_2$ can be calculated by GM1 and GM2. Hence, the matrix of Mirror 2 is

$$\mathrm{M_1} = I - 2n_1 \cdot {n_1}^T$$

$$= \begin{bmatrix} 1 - \frac{2\cos^2 2\alpha \sin^2\theta}{1-\cos^2 2\alpha\cos^2\theta} & -\frac{\sin 4\alpha \sin\theta}{1-\cos^2 2\alpha\cos^2\theta} & 0 \\ -\frac{\sin 4\alpha \sin\theta}{1-\cos^2 2\alpha\cos^2\theta} & 1 - \frac{2\sin^2 2\alpha}{1-\cos^2 2\alpha\cos^2\theta} & 0 \\ 0 & 0 & 1 \end{bmatrix}.$$

After a series of reflections, the overall effective mirror matrix is

$$\mathrm{M} = \mathrm{G_2}\mathrm{M_1}\mathrm{G_1}$$

$$= -\begin{bmatrix} -\frac{2\sin 2\alpha+\cos^2 2\alpha\cos 2\theta-2\sin 2\alpha\cos^2 2\theta+\cos^2 2\alpha\cos^2 2\theta-2\cos^2 2\theta}{\cos^2 2\alpha\cos 2\theta+\cos^2 2\alpha-2} & 0 & -\frac{\sin 2\theta(2\cos 2\theta-\cos^2 2\alpha\cos 2\theta-\cos^2 2\alpha+2\sin 2\alpha\cos 2\theta)}{\cos^2 2\alpha\cos 2\theta+\cos^2 2\alpha-2} \\ 0 & 1 & 0 \\ \frac{\sin 2\theta(2\cos 2\theta-\cos^2 2\alpha\cos 2\theta-\cos^2 2\alpha+2\sin 2\alpha\cos 2\theta)}{\cos^2 2\alpha\cos 2\theta+\cos^2 2\alpha-2} & 0 & -\frac{2\sin 2\alpha+\cos^2 2\alpha\cos 2\theta-2\sin 2\alpha\cos^2 2\theta+\cos^2 2\alpha\cos^2 2\theta-2\cos^2 2\theta}{\cos^2 2\alpha\cos 2\theta+\cos^2 2\alpha-2} \end{bmatrix},$$

M has the same structure with the rotation matrix around the y axis, which verifies the collinearity of the output images and also shows that the output images will be only rotate around y axis. When $\alpha = 0°$, we have

$$\mathrm{M}' = -\begin{bmatrix} \cos(-2\theta) & 0 & \sin(-2\theta) \\ 0 & 1 & 0 \\ -\sin(-2\theta) & 0 & \cos(-2\theta) \end{bmatrix}.$$

In the M' matrix, $-2\theta$ indicates the image is rotated by $-2\theta$, and the minus sign in front of the matrix indicates that the image is flipped in all axes. In **Supplementary Figure 3**, the y-axis represents the propagation direction of light, so the flip in y-axis means the output of the propagation direction is inverse. The flip of x- and z-axes simply indicates the output image is flipped on the x-z plane. The $-2\theta$ rotation of the image when $\alpha = 0°$ can also be confirmed in the Zemax simulation (**Supplementary Figure 5**, middle). We then find that the relative image rotation angle ($\gamma$) between two output images with $\alpha = 0°$ and $\alpha°$ is

$$\gamma = \sin^{-1}\left(-\frac{\sin 2\theta\left(2\cos 2\theta-\cos^2 2\alpha\cos 2\theta-\cos^2 2\alpha+2\sin 2\alpha\cos 2\theta\right)}{\cos^2 2\alpha\cos 2\theta+\cos^2 2\alpha-2}\right) + 2\theta.$$

In **Supplementary Figure 4**, we plot the relative image rotation angle for different choices of galvo orientation angle $\theta$ and different scan angles $\alpha$ of the glavo mirros. As one can see, through a judicious choice of the parameters, image rotators covering a wide range of rotation angles can be created.

We have also simulated our image rotator using Zemax. The resulting schematics show the 3D layout of the system, and the beam path of the galvo mirrors rotate at -10, 0, and 10 degrees. The top row in **Supplementary Figure 5** shows the isometric view and the bottom row shows the front view of the system. Two galvo mirrors are shown in the figure and there are three static mirrors for each rotation angle (only the one upon which light is incident is shown in each figure). In addition, two static mirrors below the galvo mirrors are added, which are desirable in an experimental implementation: with those two mirrors, one can keep the input and output beam collinear (i.e. traveling in the same direction). The presence of these two additional mirrors however do not affect the working principle of the rotator.

For an image rotator with three static mirrors (three image rotation angles), a light path difference between the center mirror ($\alpha$ =0) and the static mirrors above or below exists.

As shown in **Supplementary Figure 6**, the light path difference between $\overline{P_1OP_3}$ and $\overline{P_1P_2P_3}$ is

$$\Delta l = 2l(\frac{1}{\cos(2\alpha)} - 1),$$

where $l = \overline{P_1O}$. $M_1$ and $M_2$ represent two static mirrors between two galvo mirrors ($G_1$ and $G_2$).

In practice in our OPSIM, the light path difference will introduce a shift in the z-direction (along the optical axis of the primary objective) to the light-sheet and the detection focal plane. Thus, while the light-sheet and detection focal plane stay in alignment, the resulting data is slightly shifted (i.e. re-focused) in z in respect to the other two directions. In our OPSIM system, the resulting z-offset is on the order of 400nm, which is corrected in the data preprocessing steps when the three directions are registered to each other.

Importantly, for two static mirrors arranged symmetrically above and below the galvo plane, no path length difference occurs between the two rotated images. I.e., an image rotator with two rotation angles, whose magnitude can be selected in the design (e.g. + 45 and -45 degrees), would feature the same path length for both arms. In contrast, a path length difference occurs once three or more rotation angles are implemented.

#### Extended Figures

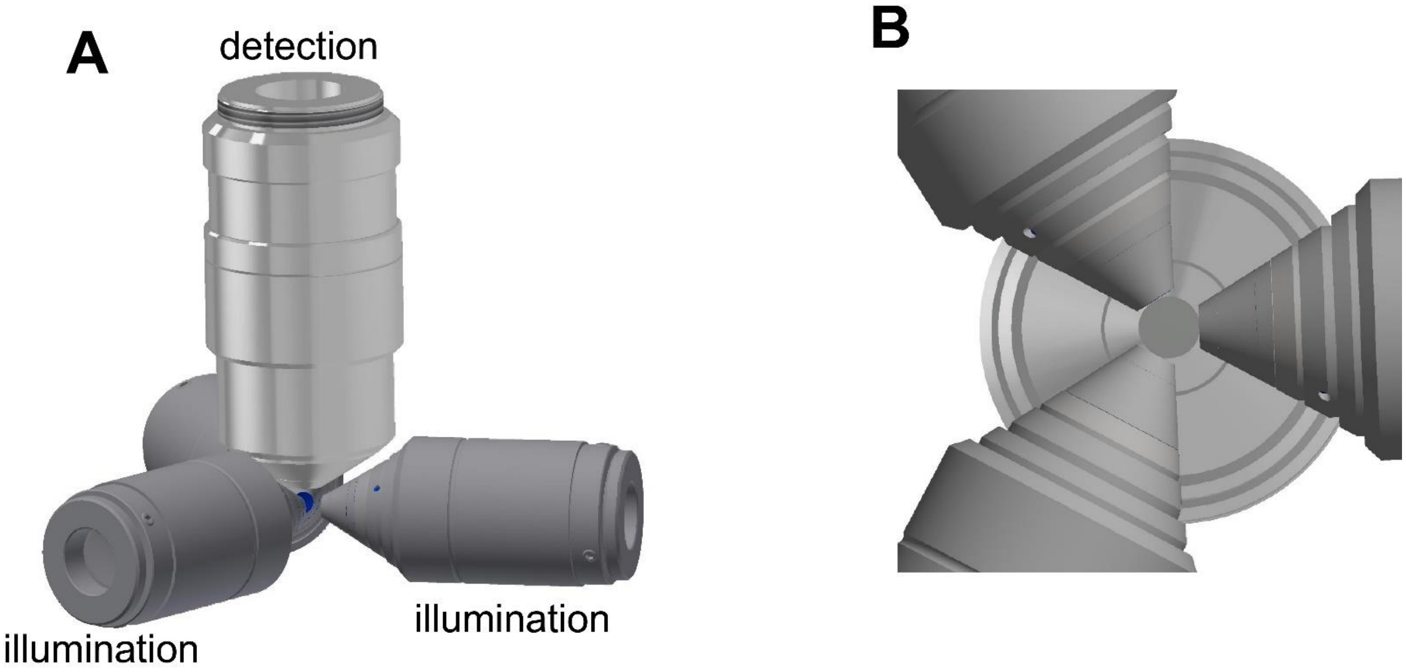


**Extended Figure 1 – Conceptual setup for structured illumination with three illumination directions. A** Rendering of a potential setup for multi-directional LSFM-SIM, consisting of three illumination objectives, angled at 120 degrees to each other, and one detection objective. **B** Bottom view of the objective assembly.

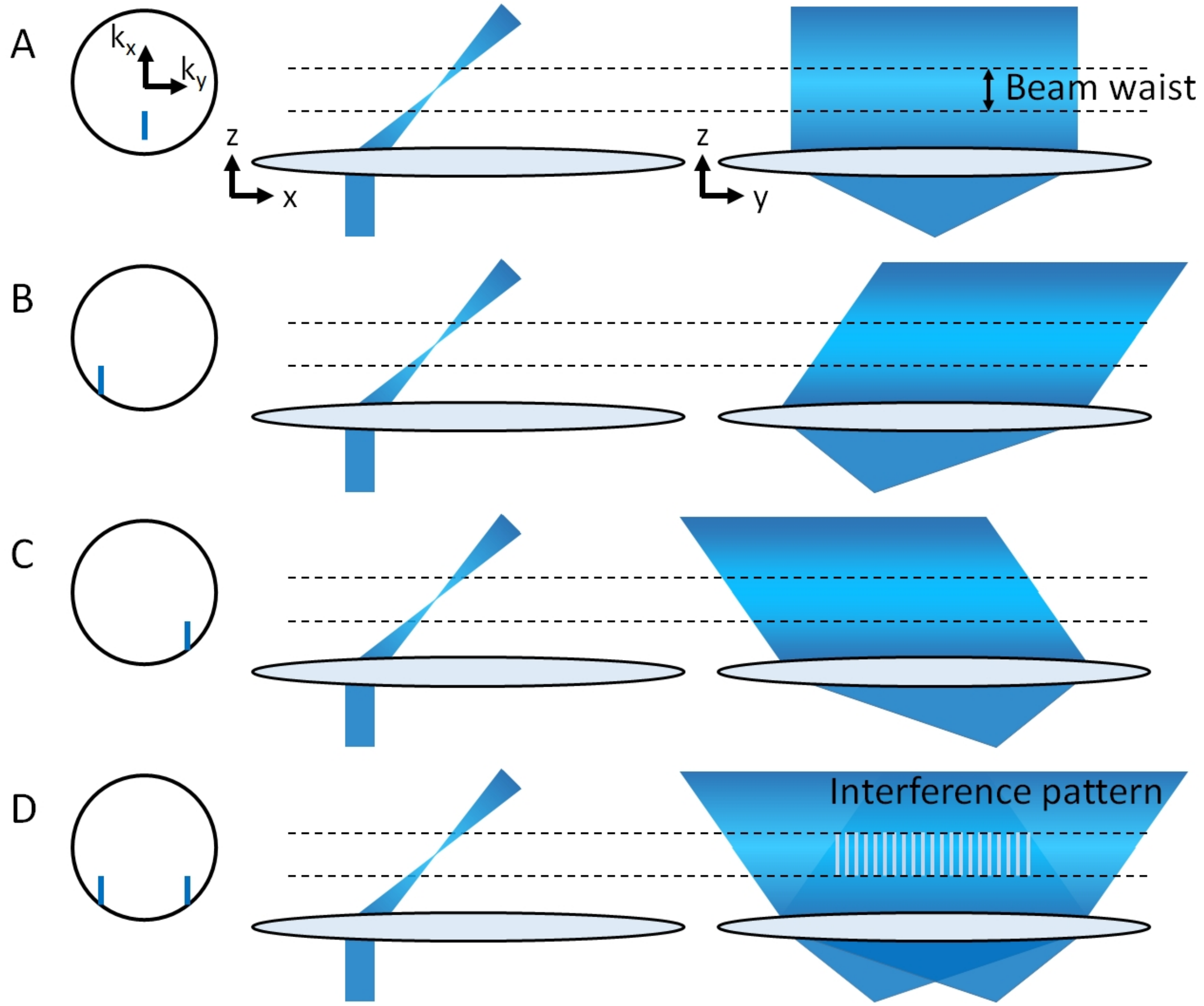


**Extended Figure 2 –Schematic illustration of the oblique plane structured illumination sheet. A:** Electric field (blue stripe) in the pupil of the primary objective in an OPM system. The offset in the negative $k_x$ direction of the electric field (i.e. the blue stripe is not centered in the pupil) causes the light-sheet to be tilted in sample space. Middle: x-z view of the oblique light-sheet intensity distribution as it emerges from the objective. Right: y-z view of the light-sheet intensity distribution. **B** The electric field in the pupil has been shifted to the left (in the minus $k_y$ direction). As a result, in the y-z view, the light-sheet is angled and propagates diagonally (in a positive y direction). **C** The electric field in the pupil has been shifted to the right (positive $k_y$ direction). As a result, in the y-z view, the light-sheet is angled and propagates diagonally (in a negative y direction). **D** Coherent superposition of the electric fields in **B-C** results in a 1D interference pattern along the y direction as seen in the y-z view. Importantly, even though the individual light-sheets are angled (as viewed in an y-z plane), the beam waist of each sheet remains parallel to the focal plane of the primary objective. The dotted lines in **A-D** indicate the limits of the beam waist. In other words, the beam waists are not rotated in an y-z view, but rather form a parallelogram.

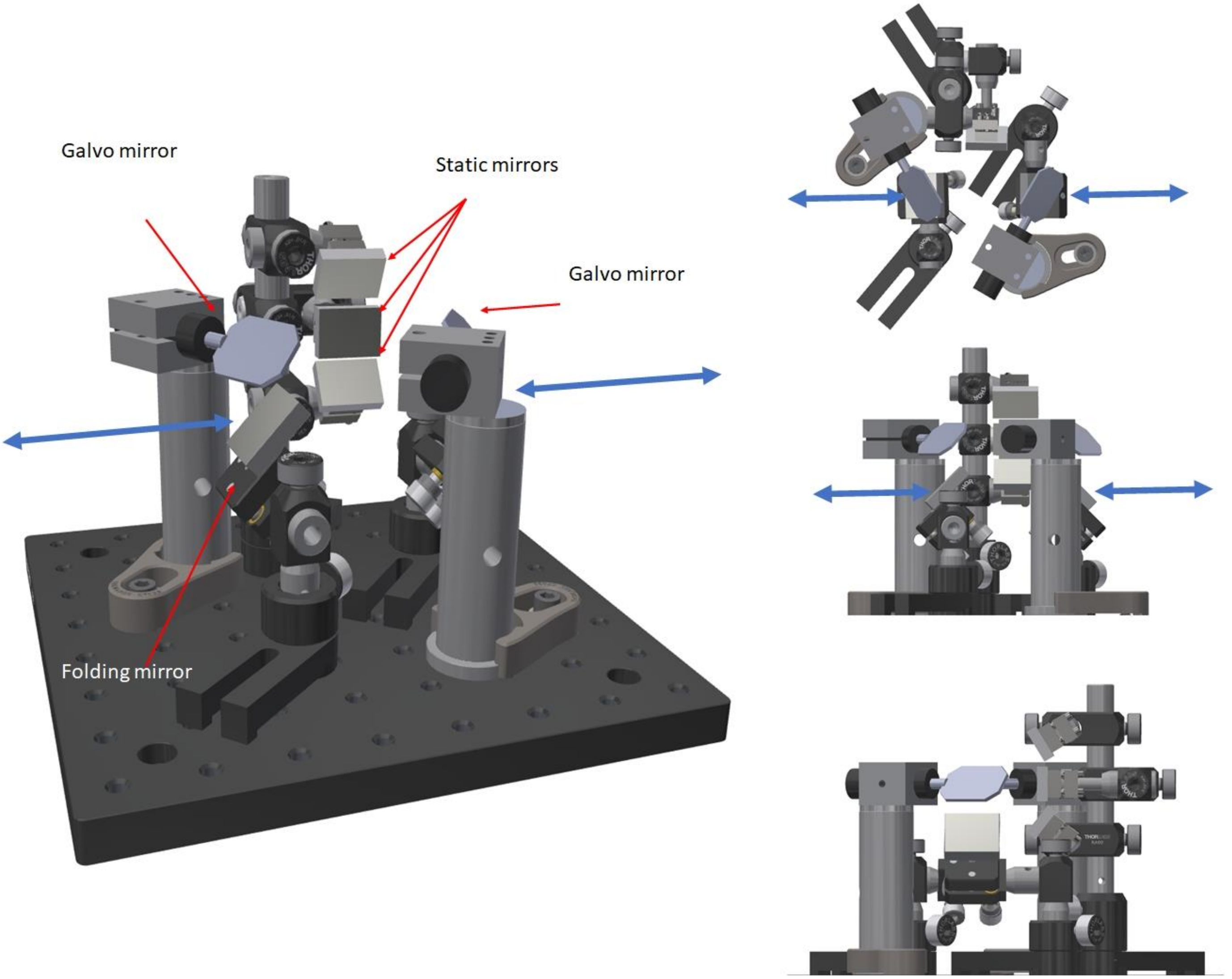


**Extended Figure 3** Rendering of the image rotator unit (left) and a top, side and frontal view (right). The main components are two galvo mirrors and three static mirrors. Two folding mirrors bring the in and output beam (blue arrows) on a common optical axis

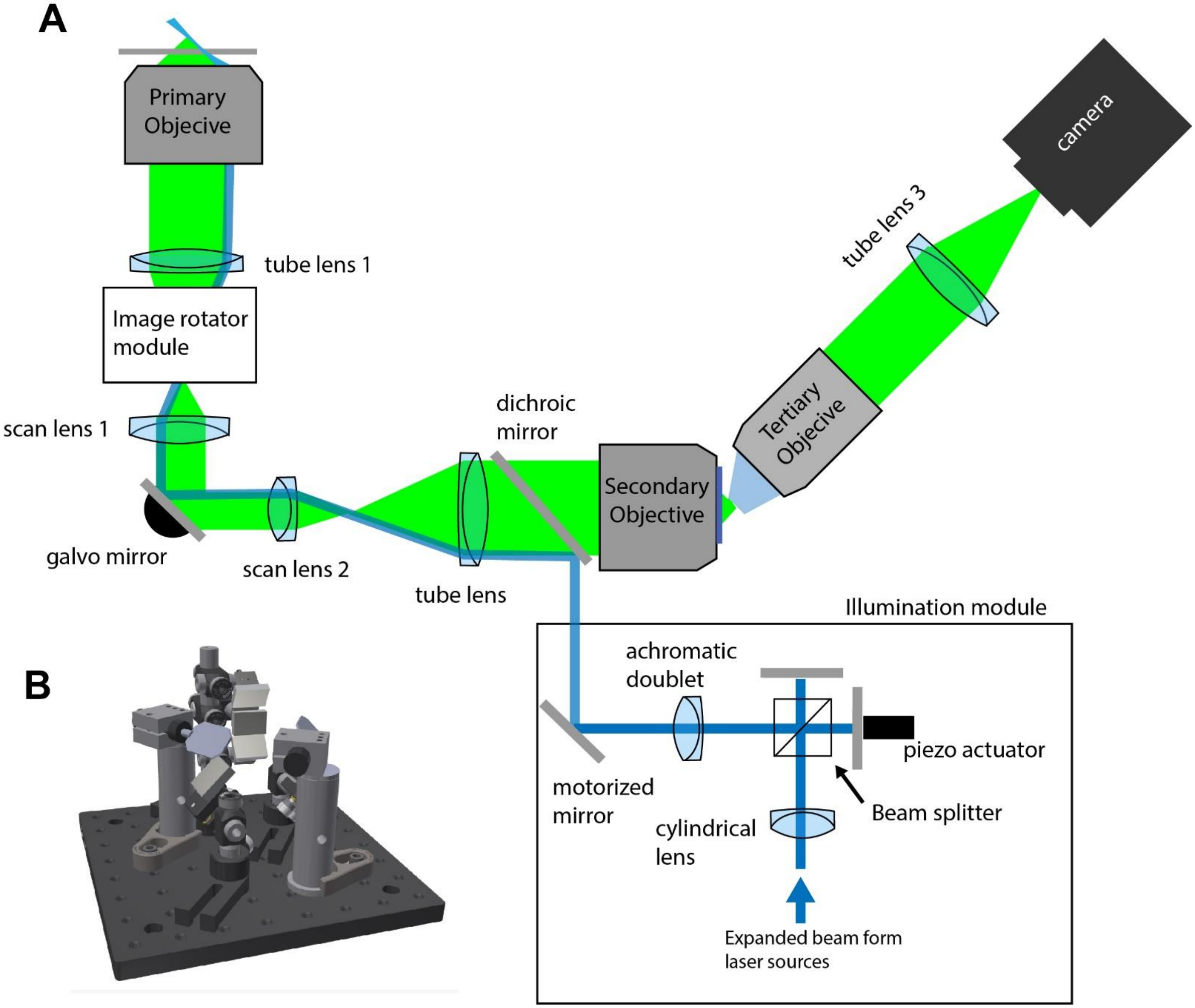


**Extended Figure 4 – Setup for Oblique Plane Structured Illumination Microscopy. A** Schematic drawing of the setup. The illumination unit creates to interfering light-sheets (blue), which can be phase stepped by a piezo actuator. They are coupled into the OPSIM microscope with a dichroic mirror. The image rotator module rotates the light-sheets by three discrete steps. The returning fluorescence light (green) is de-rotated by the image rotator to align with the alignment of the tertiary objective.

**B** Rendering of the image rotator module. Two galvanometric mirrors are used to select three beam paths, each of which imparts a different amount of image rotation.

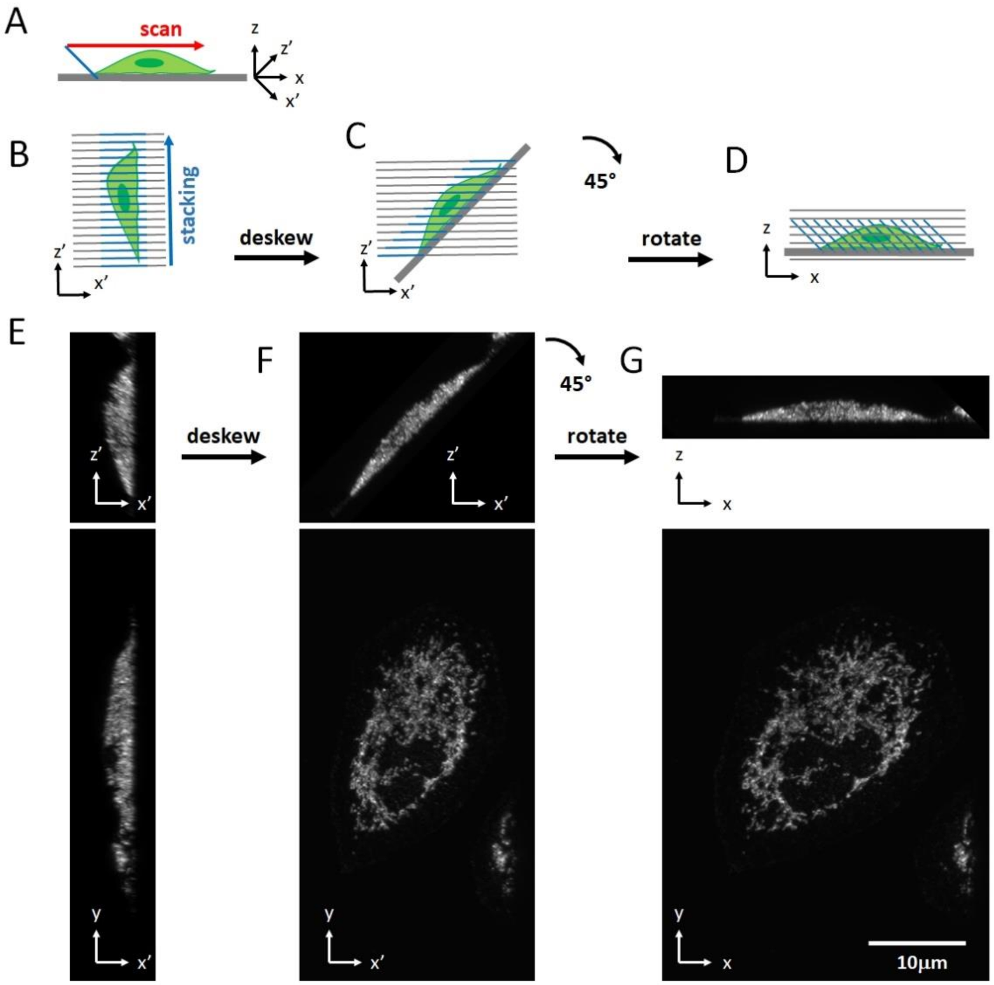


**Extended Figure 5 –Workflow for data preprocessing, part 1.** Prior to SIM processing, the data is pre-processed: Each raw stack is de-skewed and rotated into the coverslip (x-y-z) reference frame as it is schematically shown in **A-D** and on biological data from **E-F.** **A** To scan a volume, the light-sheet and detection focal plane (blue line) are scanned along the coverslip. x-z is the coordinate system of the primary objective, with z along its optical axis. x'-z' is the coordinate frame of the light-sheet and the tilted detection focal plane. **B** The images from a scan are assembled into a stack. As the scan direction is not along the z' axis, the resulting stack is skewed (i.e. each plane has a desired displacement in z', but also an unwanted one in x'). **C** De-skewing results in a proper x'-z' stack. **D** The data is rotated into a x-z reference frame. Empty data, introduced by the de-skewing, above the cell and below the coverslip is discarded. **E-F**: maximum intensity projections of an U2OS cell labeled for MIC60 at different stages of the pre-processing.

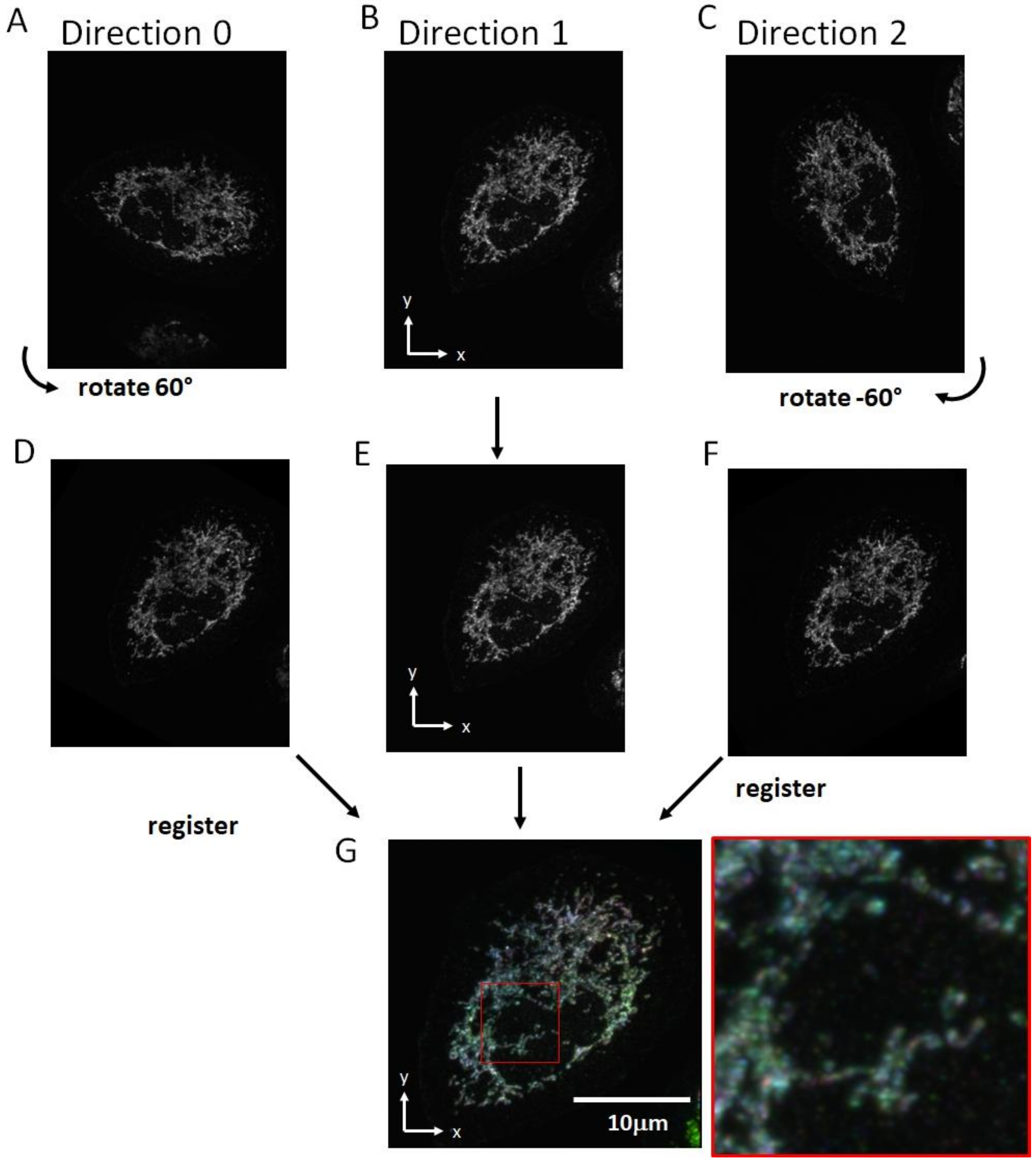


**Extended Figure 6 –Workflow for data preprocessing, part 2. A-F** After de-skewing and rotating the data into an x-y-z coordinate frame of the primary objective/coverslip, the data volumes for the first (**A**, Direction 0) and the third (**C,** Direction 2) are rotated by plus and minus 60 degrees around the z- axis to share the same orientation as the second direction (**B**, Direction 1). **G** The rotated data sets for Direction 0 and Direction 1 are then registered to Direction 1, which have been color coded here for clarity. Inset shows a magnified version of the red boxed region.

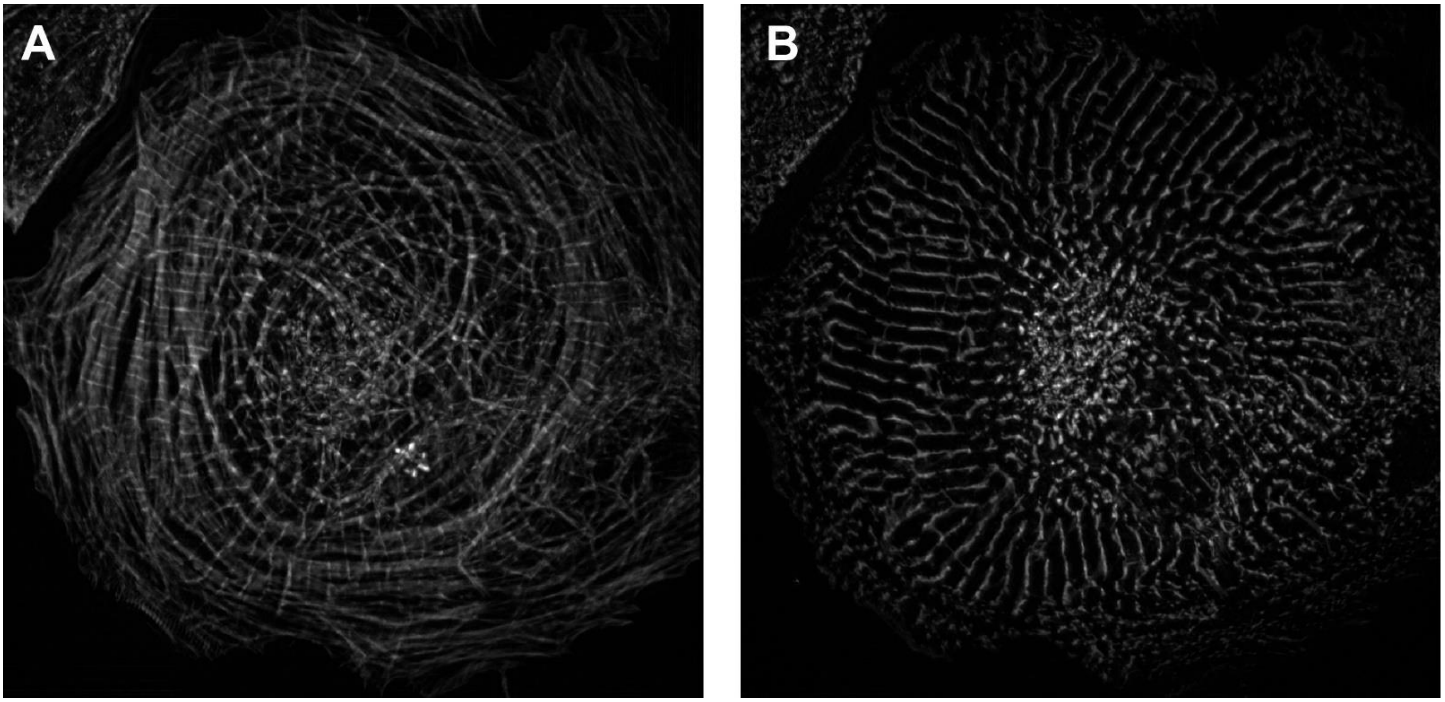


**Extended Figure 7 – Cardiomyocyte labeled with Phalloidin and Actinin. A** Phalloidin labeled channel of the cell shown in **Figure 3 C** as imaged by OPSIM. **B** Alpha-Actinin 2, labeled with Alexa 561, as imaged by OPSIM.

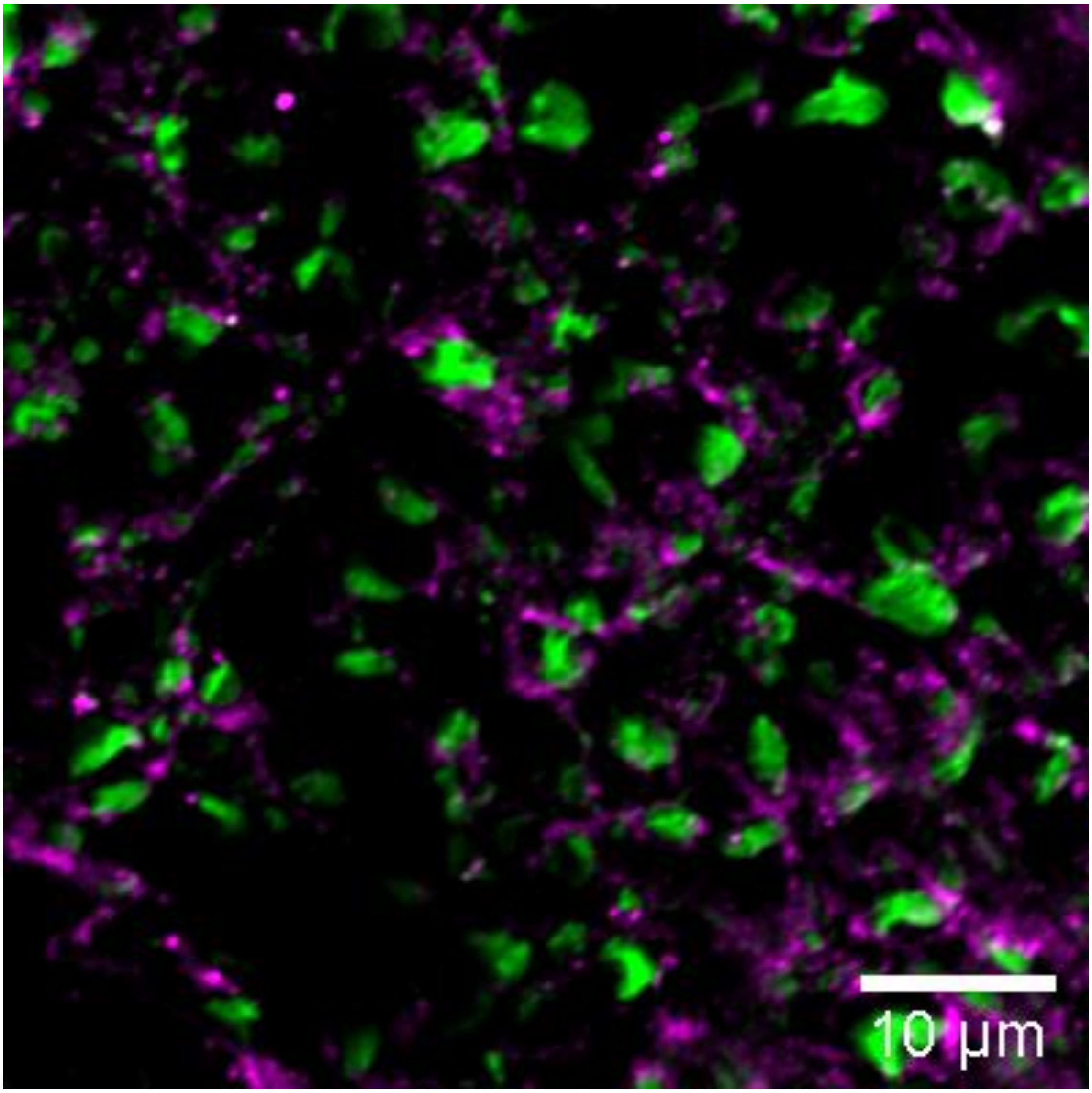


**Extended Figure 8 – spinal cord slice imaged with a Confocal microscope.** Single plane of neurofilament (NF200, green) and myelin (PLP, magenta) in a 20 micron thick spinal cord slice, as imaged with a Confocal microscope.

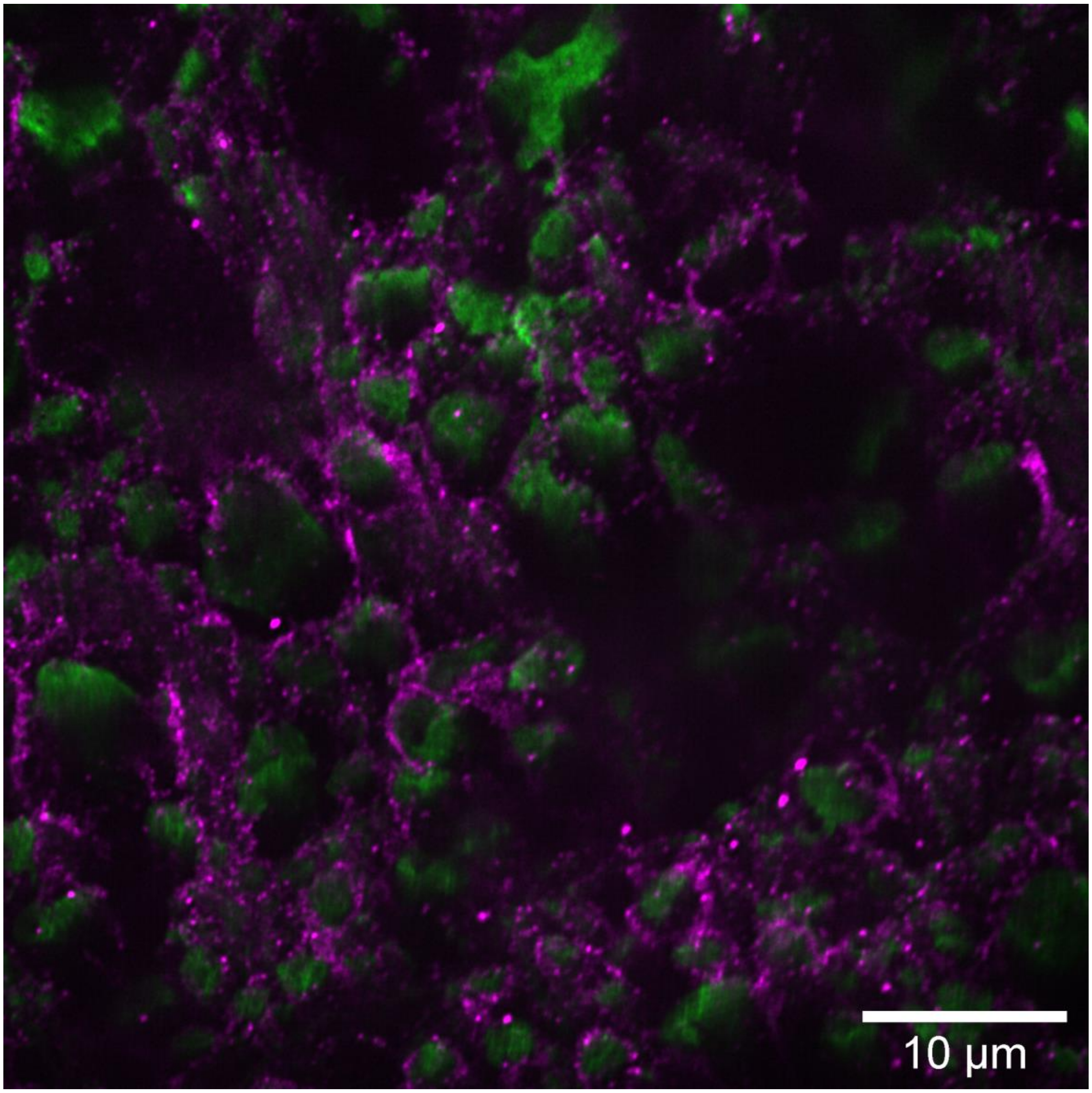


**Extended Figure 9 – spinal cord slices imaged with a Nikon SoRa spinning disk .** Single plane of neurofilament (NF200, green) and myelin (PLP, magenta) in a 20 micron thick spinal cord slice, as imaged with a SoRa spinning disk.

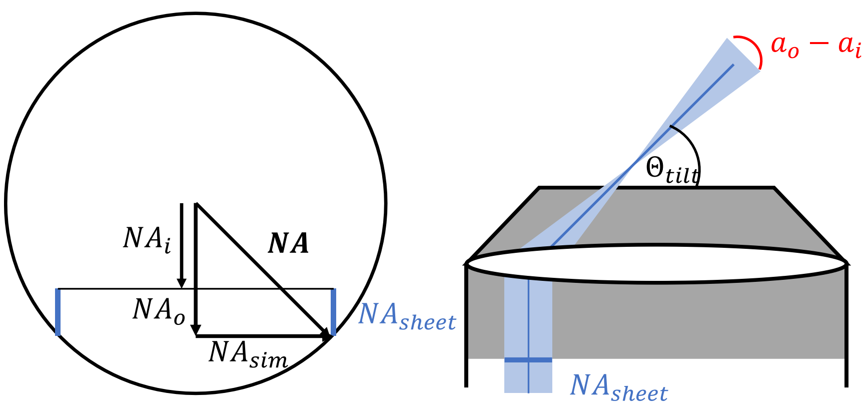


$$NA = n\sin(a_{max})$$
$$NA_o = n\sin(a_o)$$
$$NA_i = n\sin(a_i)$$
$$NA_{sim} = \sqrt{NA^2 - {NA_o}^2}$$

***Conditions***:

$NA = 1.35\ \&\ n = 1.4$

***If***

$\Theta_{tilt} = 45°$,

$$NA_{sheet} = n\sin\left(\frac{a_o - a_i}{2}\right) = 0.12$$

***Then***

$a_o - a_i = 10°$, $a_o = 50°$

$\boldsymbol{NA_{sim} = 0.82}$

**Extended Figure 10 –Excitation intensity in the primary pupil in Oblique Plane Structured Illumination Microscopy.** Schematic drawing of the pupil of the primary objective (numerical aperture: 1.35) in OPSIM, and the location of the two thin stripes (blue) that create the structured, oblique light-sheet. For a tilt angle of 45 degrees, a light-sheet NA of ~0.12, an NA for structured illumination of 0.82 results. Experimentally, the highest excitation NA we have achieved was 0.79, before running into beam clipping or vignetting effects. For conventional SIM, the highest NA that can be used for pattern generation equals to the NA of the primary objective, which illustrates the tradeoff that needs to be done for the tilted structured light-sheet.

#### Supplementary Figures

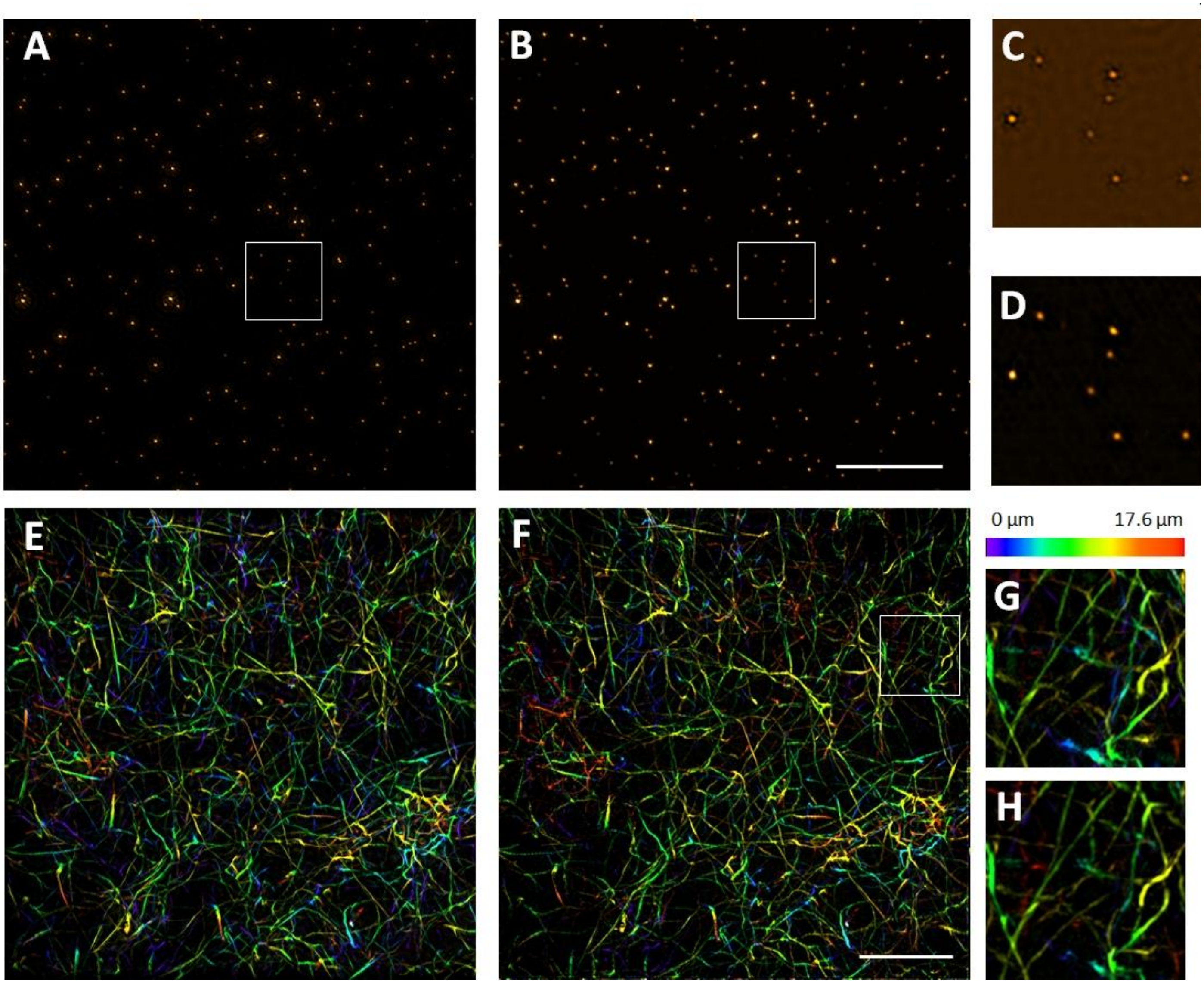


**Supplementary Figure 1 –Comparison of SIM reconstructions. A-D** 100nm fluorescent nanospheres as imaged by OPSIM. **A** SIM reconstruction with Wiener deconvolution using an experimental PSF. Maximum intensity projection. **B** SIM reconstruction with Richardson Lucy deconvolution using a synthetic PSF. Maximum intensity projection. **C** Magnified boxed region shown in **A**. Single image plane. **D** Magnified boxed region shown in **B**. Single Image plane. **E-H** Fluorescence collagen sample, as imaged by OPSIM. Color encodes height above the coverslip. **E** SIM Reconstruction with Wiener deconvolution using an experimental PSF. Maximum Intensity Projection. **F** SIM Reconstruction with Richardson Lucy deconvolution using a synthetic PSF. Maximum Intensity Projection **G** Magnified boxed region shown in **E**. Maximum Intensity Projection. **H** Magnified boxed region shown in **F**. Maximum Intensity Projection. The displayed intensities range from smallest to largest value in each image. Undershoots of conventional Wiener deconvolution are not visible in **A,E,G** as the images are shown as maximum intensity projections. Scale Bar: 10 microns.

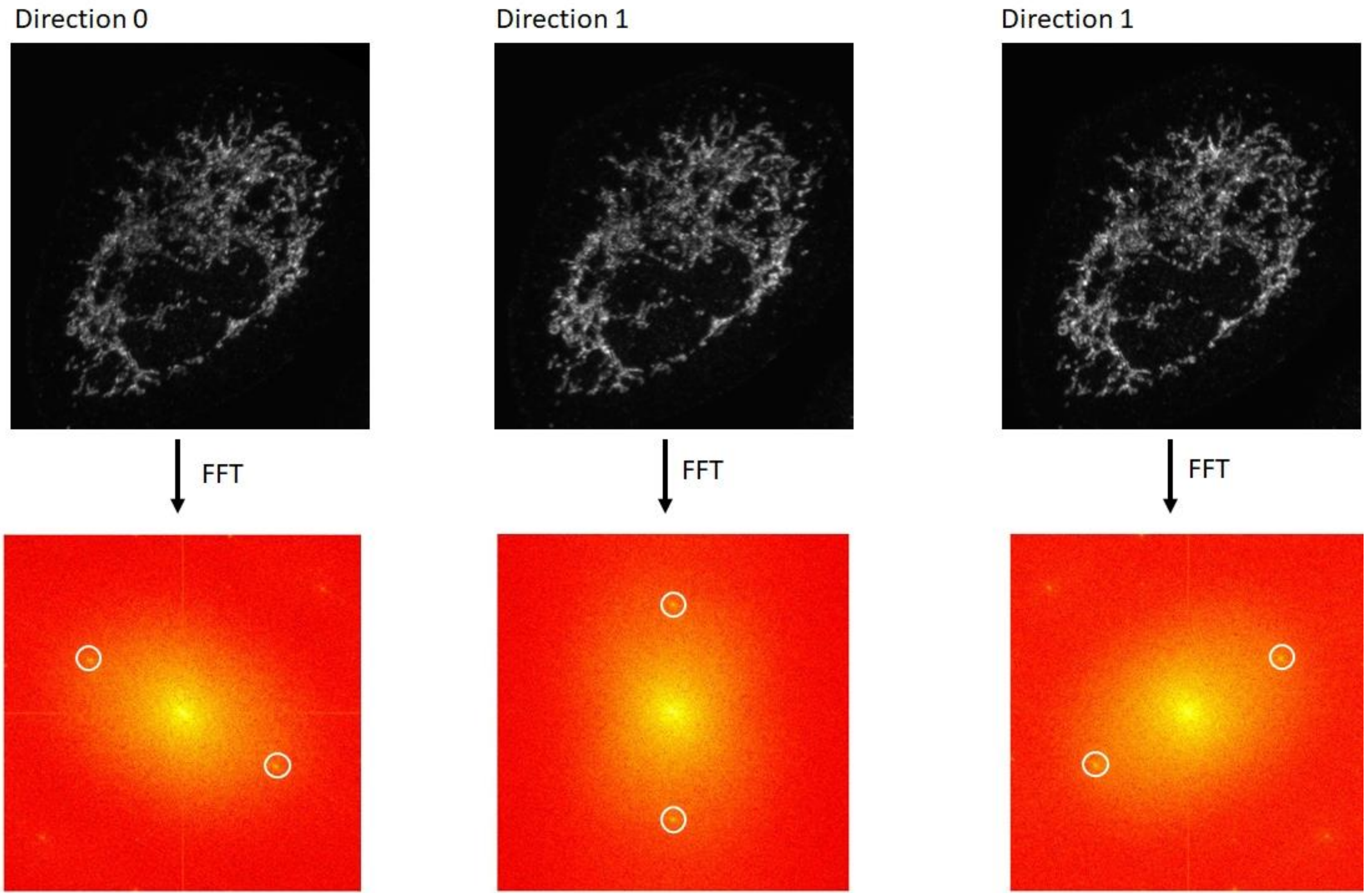


**Supplementary Figure 2 – Fourier transforms of OPSIM raw data.** Raw OPSIM data for the three directions after pre-processing. Top row shows maximum intensity of a MIC60 labelled U2OS cell shown in **Figure 2I**. Bottom row shows the Fourier transforms of the images above, shown as the log of a power spectrum. White circles highlight the peaks of the structured illumination pattern.

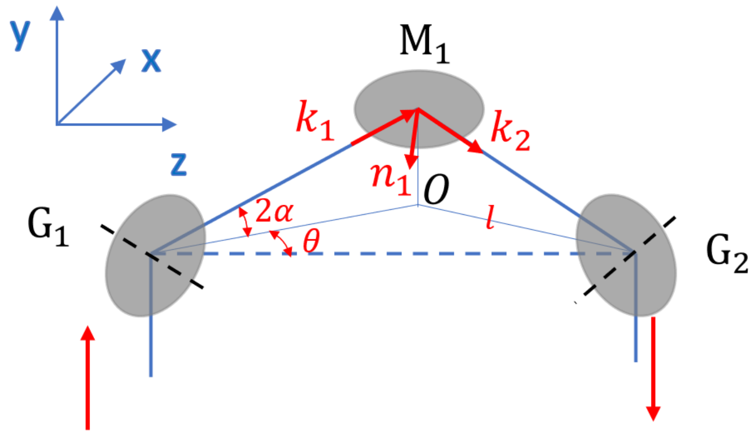


**Supplementary Figure 3** Illustration of the image rotator. The galvos are rotated relative to the X-axis by $\pm\theta$. The galvo mirror both rotate by the angle $\pm\alpha$, resulting in a deflection of the light by $\pm 2\alpha$.

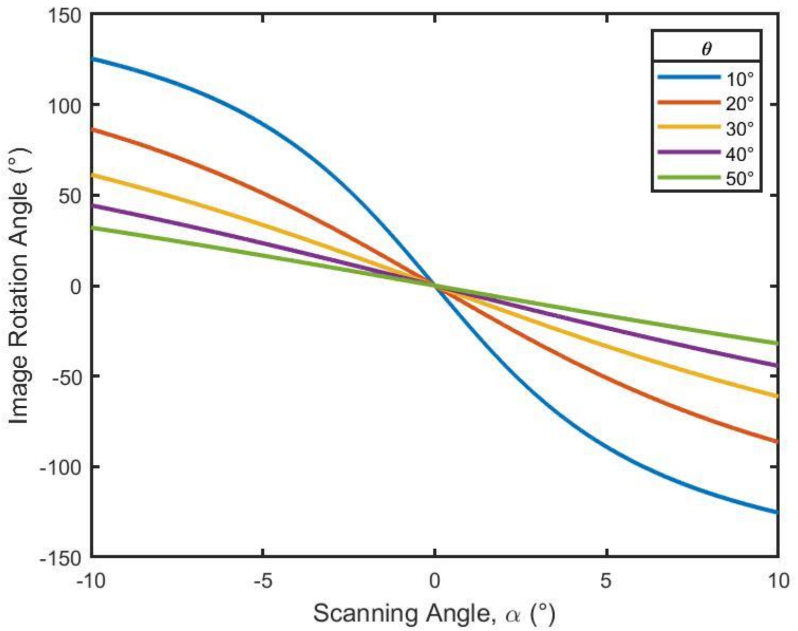


**Supplementary Figure 4** The relative image rotation angle ($\gamma$) versus Galvo scanning angle ($\alpha$).

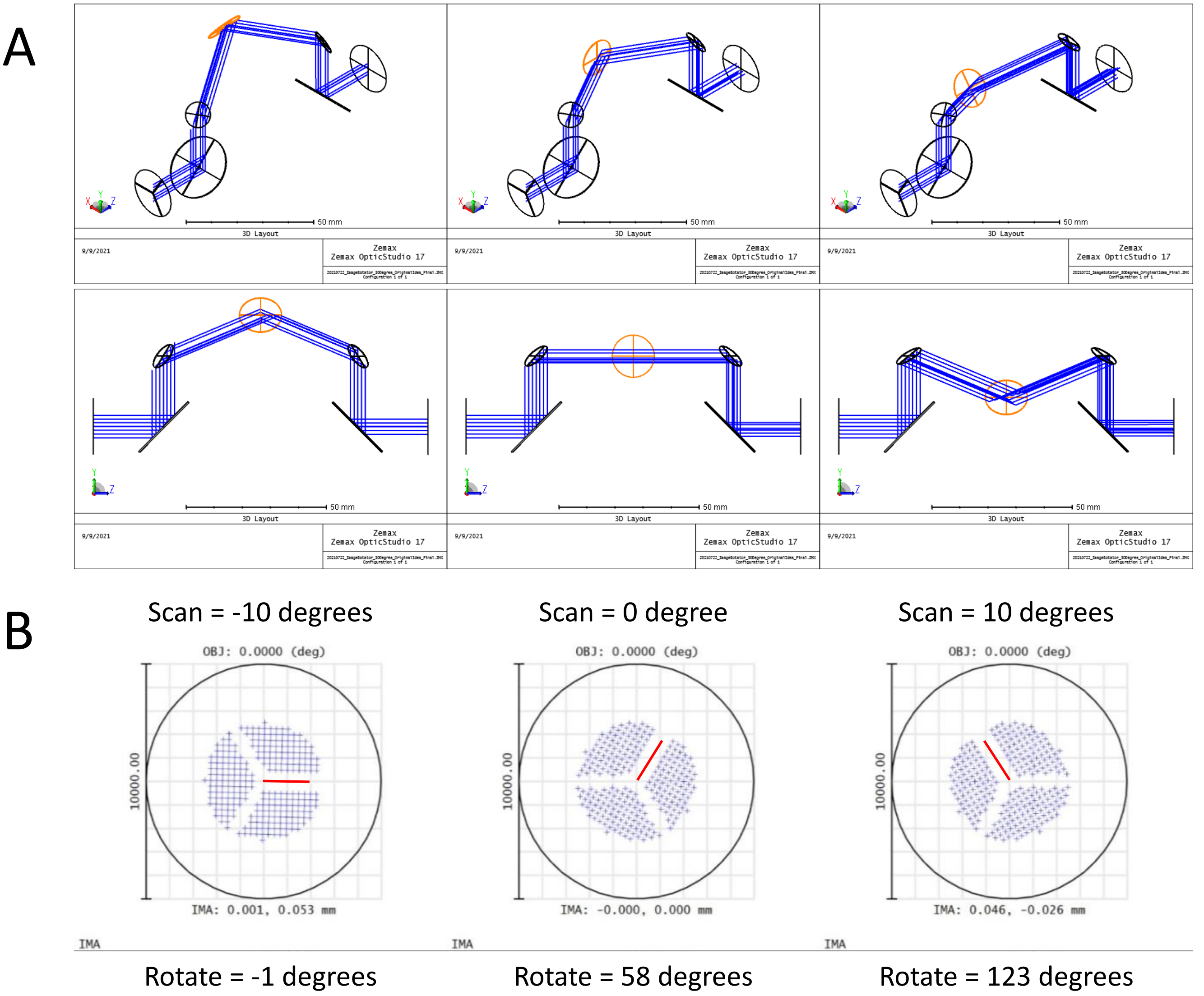


**Supplementary Figure 5** Zemax simulation of the image rotator. In this simulation, $\theta = 30^{\circ}$. **A** The static mirrors (only one shown at a time) are shown in orange. **B** The rotation of the image simulated with Zemax. Image can be rotated over a range of ~124 degrees for mechanical galvo mirrors rotations of -10 and 10 degrees, respectively.

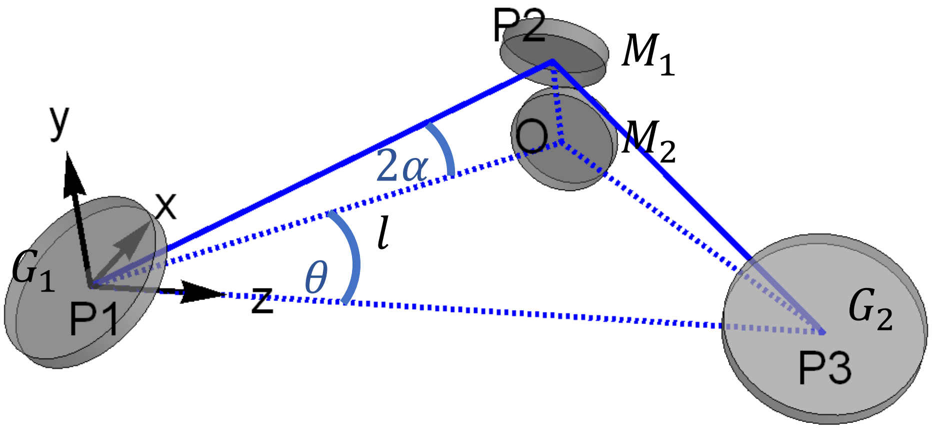


**Supplementary Figure 6** Calculation of the optical path difference between the central static mirror and one of the outer static mirrors for the setup used in this manuscript allowing three image rotation angles.

#### Supplementary Tables

| Microscope | FWHM x | FWHM y | FWHM z |
|---|---|---|---|
| **OPSIM** | 148 ± 10nm | 138 ± 12nm | 493 ± 51nm |
| **LLSM (n=1)** | 280nm | 147nm | 388nm |
| **SoRa** | 134 ± 7nm | 128 ± 5nm | 429. ± 3nm |
| **iSIM** | 160 ± 21nm | 166 ± 9nm | 546 ± 65nm |

**Supplementary Table 1 – Full width half maximum measurements for the microscope systems compared in this manuscript.**

| Figure | microscope | Exposure time | Acquisition rate | Voxel size (x,y,z) | Laser power | Post processing |
|---|---|---|---|---|---|---|
| **1G,I** | **OPSIM** | **1ms** | **1.03Hz** | **57x57x114nm** | **0.616mW** | **SIM reconstruction** |
| **2A-C** | **OPSIM** | **10ms** | **---** | **57x57x114nm** | **50µW** | **SIM reconstruction** |
| **2D** | **LLSM SIM** | **10ms** | **--** | **69x69x89nm** | **Not recorded** | **SIM reconstruction** |
| **2E** | **SoRa** | **100ms** | **--** | **23.2x23,2x200nm** | **915µW** | **Proprietary deconvolution in Nikon software** |
| **2F** | **iSIM** | **1000ms** | **--** | **43x43x100nm** | **1mW 488nm 0.37mW 561nm** | **Deconvolution (ImageJ plugin "Microvolution")** |
| **2G** | **OPSIM** | **20ms** | **--** | **57x57x114nm** | **128µW 488nm 30µW 561nm** | **SIM reconstruction** |
| **2H** | **iSIM** | **500ms** | **--** | **43x43x200nm** | **0.79mW 488nm 0.74mW 561nm** | **Deconvolution (ImageJ plugin "Microvolution")** |
| **2I** | **OPSIM** | **10ms** | **--** | **57x57x114nm** | **128µW** | **SIM reconstruction** |
| **2K** | **SoRa** | **226ms** | **0.083Hz** | **23.2x23,2x200nm** | **935µW** | **Proprietary deconvolution in Nikon software** |
| **2L** | **OPSIM** | **8ms** | **0.132Hz** | **57x57x114nm** | **50µW** | **SIM reconstruction** |
| **3A** | **OPSIM** | **10ms** | **0.0660Hz** | **57x57x114nm** | **175µW** | **SIM reconstruction** |
| **3B** | **OPSIM** | **5ms** | **0.1754Hz** | **57x57x114nm** | **175µW** | **SIM reconstruction** |
| **3C-D** | **OPSIM** | **10ms** | **--** | **57x57x114nm** | **50µW 488nm 46µW 561nm** | **SIM reconstruction** |
| **3E** | **OPSIM** | **25ms** | **--** | **57x57x114nm** | **30µW 488nm 63µW 561nm** | **SIM reconstruction** |
| **4A-D** | **OPSIM** | **1ms** | **0.8619Hz** | **57x57x114nm** | **0.616mW** | **SIM reconstruction** |
| **4E-G** | **OPSIM** | **0.76ms** | **1.2232Hz** | **57x57x114nm** | **0.616mW** | **SIM reconstruction** |
| **Movie 1** | **OPSIM** | **1ms** | **0.8619Hz** | **57x57x114nm** | **0.616mW** | **SIM reconstruction** |
| **Movie 2** | **OPSIM** | **0.76ms** | **1.2232Hz** | **57x57x114nm** | **0.616mW** | **SIM reconstruction rolling average over one frame** |
| **Movie 3** | **OPSIM** | **1ms** | **1.4289Hz** | **57x57x114nm** | **0.616mW** | **SIM reconstruction** |

**Supplementary Table 2 – Image acquisition parameters** Laser power was measured in the pupil of the objective. For a single timepoint acquisition, no acquisition rate is reported. All OPSIM reconstructions were performed with the algorithm that employs Richardson Lucy deconvolution (see also **Methods**). Rolling frame average for Movie 2 was computed over two subsequent frames.

**Supplementary Movies**

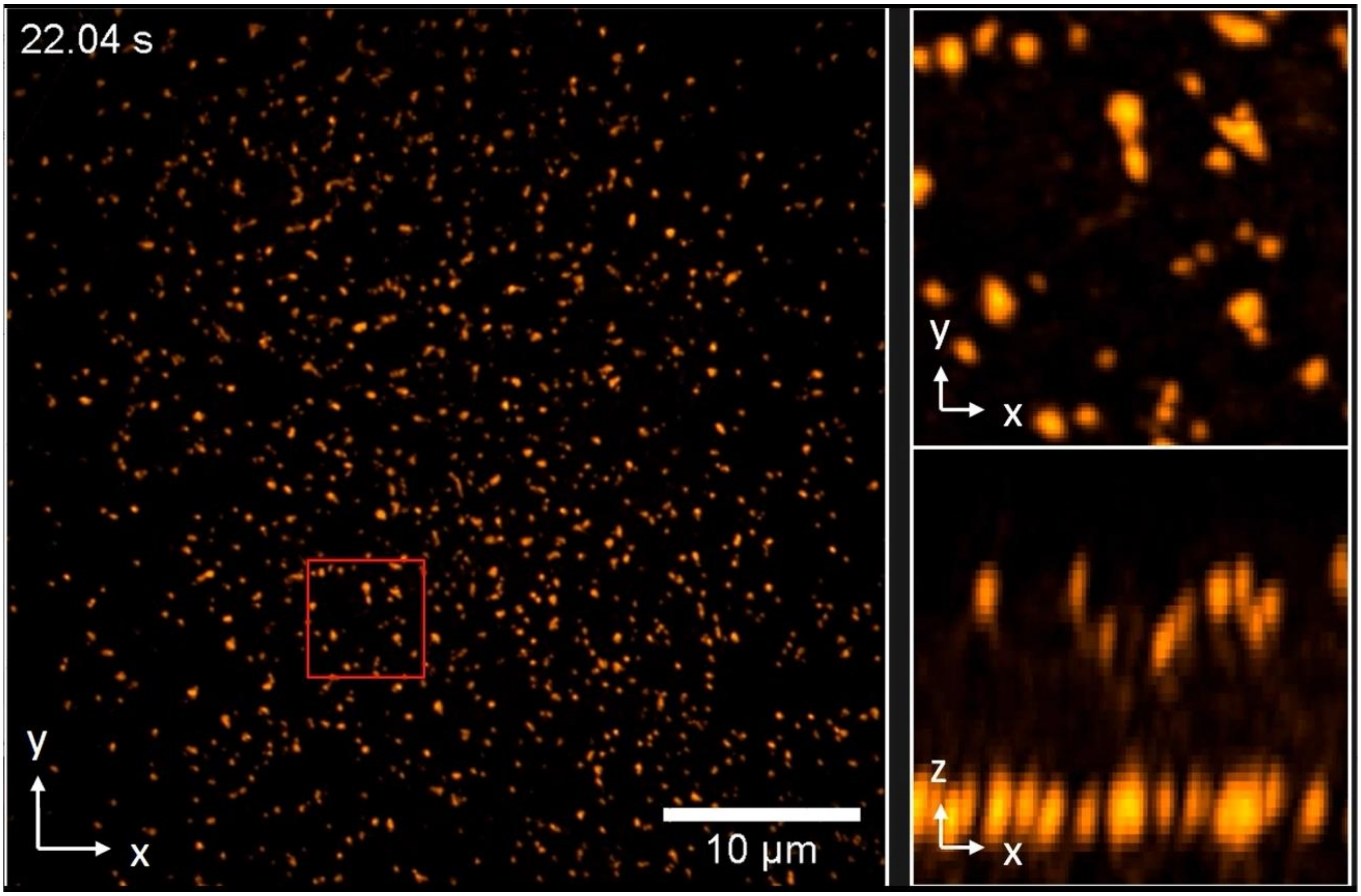


**Supplementary Movie 1 Clathrin coated vesicle dynamics.** An ARPE-19 cell, labeled for AP2-eGFP, imaged by OPSIM over 40 timepoints at a volumetric acquisition rate of 0.86 Hz (1.16s acquisition time for a full OPSIM data set for one timepoint). The left side shows a maximum intensity projection of the whole field of view, and magnified versions of the boxed regions are shown on the right.

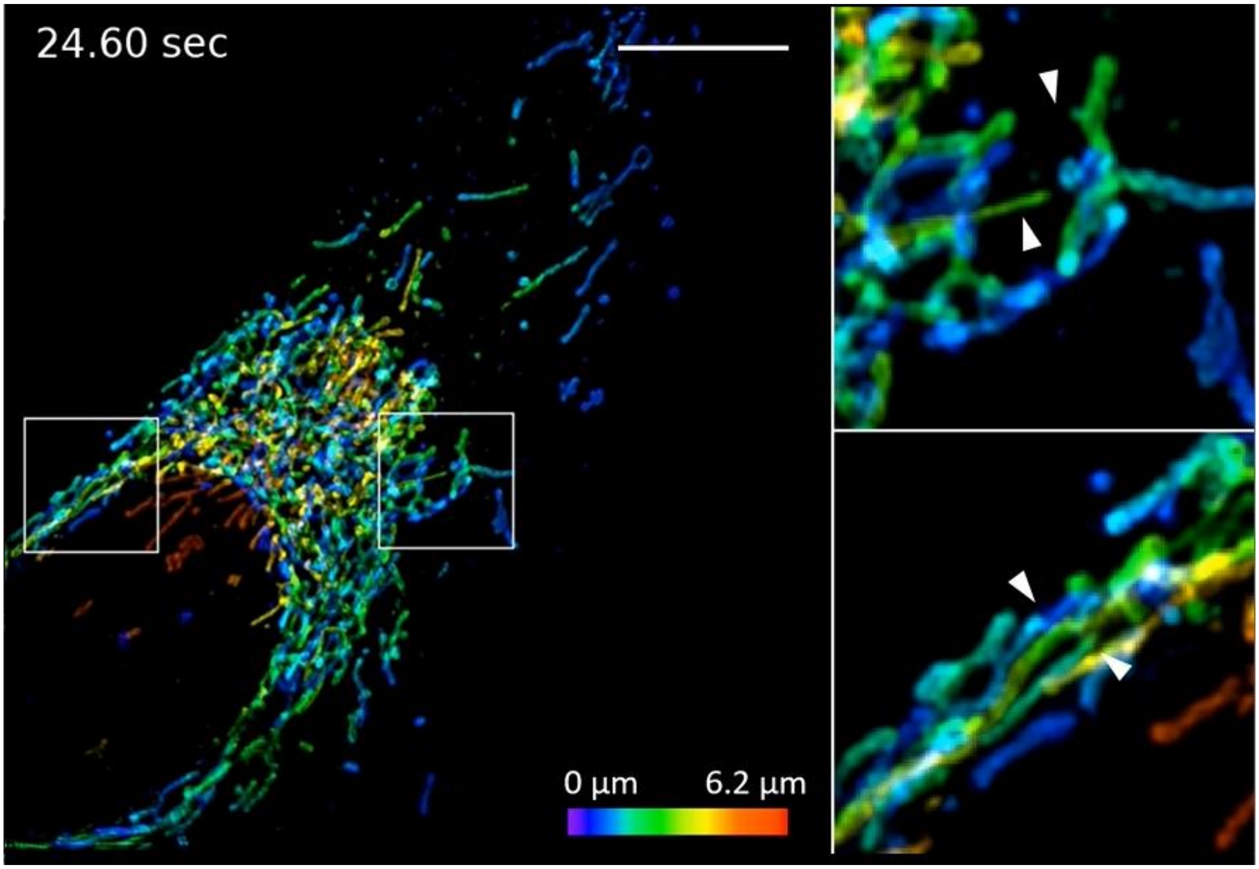


**Supplementary Movie 2 Mitochondria dynamics.** An U2OS cell, labeled for GFPOMP25, imaged by OPSIM over 38 timepoints at a volumetric acquisition rate of 1.2 Hz (0.82s acquisition time for a full OPSIM data set for one timepoint). Left shows a maximum intensity projection of the whole field of view, color coded for height. On the right, magnified versions of the boxed regions on the left are shown. White arrows point at protruding and retracting mitochondria.

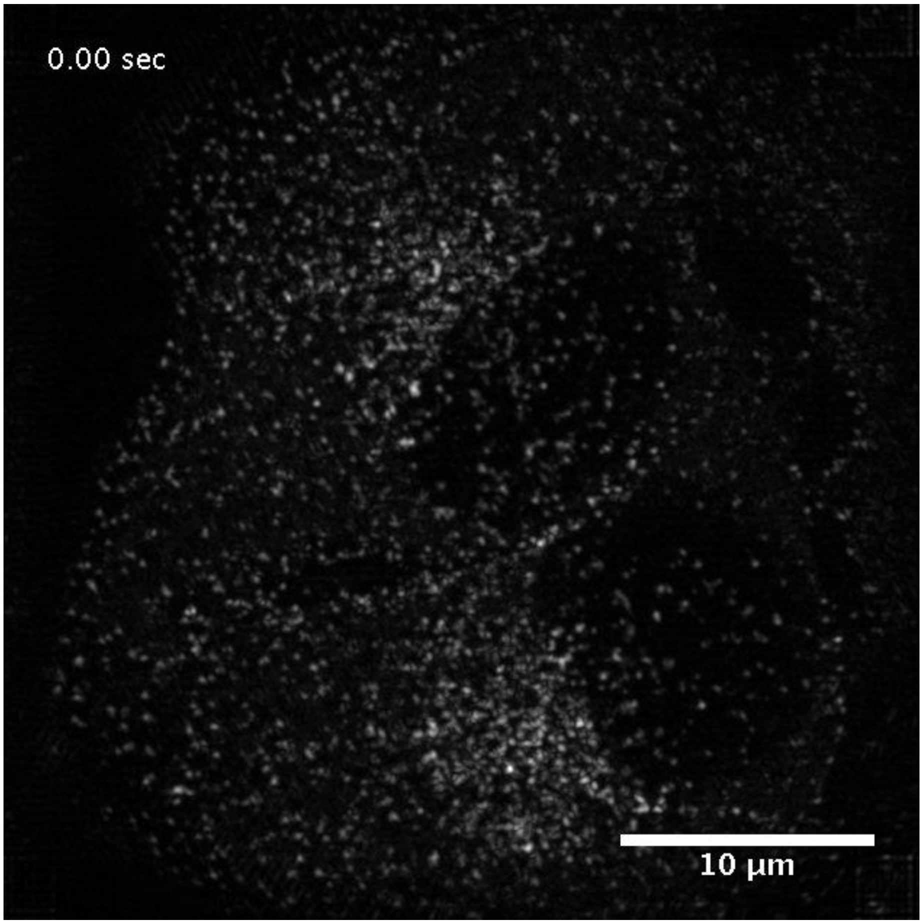


**Supplementary Movie 3 Clathrin coated vesicle dynamics.** An ARPE-19 cell, labeled for eGFP AP2, imaged by OPSIM over 38 timepoints at a volumetric acquisition rate of 1.4 Hz (0.7s acquisition time for a full OPSIM data set for one timepoint). The movie is displayed as a maximum intensity projection.
